# Jasmonate-induced prey response in the carnivorous plant *Drosera capensis*

**DOI:** 10.1101/2025.07.18.665637

**Authors:** Zane G. Long, Gemma R. Takahashi, Franchesca M. Cumpio, Omar J. Akbari, Ulysses Castelan, Mark Hadadian, Jonathan V. Le, Aden M. Alemayhu, David E. Einstein, Elliott E. Einstein, Jessica I. Kelz, Ashley O. Kwok, Allison Pineda, Pauniz Shabakesaz, Megha H. Unhelkar, Sofiya M. Woodcock, Carter T. Butts, Rachel W. Martin

**Author notes:** These authors contributed equally to this work (Zane G. Long, Gemma R. Takahashi).

## Abstract

*Drosera capensis* is a carnivorous plant native to South Africa. Central to its prey capture and digestive processes is a complex array of biochemical processes triggering the production of both enzymes and small molecules. These processes are in part activated by the release of jasmonic acid, a plant defense hormone repurposed as a prey detection signal. Here, we use RNASeq and untargeted LC-MS metabolomics to study the response of *D. capensis* to a feeding stimulus. We confirm the expression of digestive proteins predicted in prior genomic work and show up- and downregulation for a number of enzyme classes in response to jasmonic acid. Metabolomics experiments indicate that many small molecules produced during feeding depend on specific nutrient inputs from prey (and not merely a jasmonic acid stimulus). These results shed light on the molecular basis of plant carnivory and the recruitment of existing biochemical pathways to perform specialized functions.

---

Understanding how plants use molecular signals to respond to environmental stresses, and how these responses manifest in the interaction of enzymatic regulation and production of small molecules, is a key question for both basic plant biology and for the harnessing of plant biosynthesis for biotechnology applications. The jasmonic acid pathway mediates plant responses to stress, including drought, exposure to chemical cues such as fungal oligosaccharides, and wounding by herbivores (*1–4*). In response to mechanical damage, plants produce a variety of defensive compounds, including chitinases (*5,6*), protease inhibitors that disrupt insect feeding (*7–9*), as well as an array of secondary metabolites, including terpenoids (*10–13*) and flavonoids (*14–16*). Flowering plants produce large numbers of terpenoids, many of them unique to particular lineages and representing specialized attractants for beneficial organisms or toxic defenses against pests. Flavonoids and their derivatives are antioxidants whose activity is important in plant response to abiotic stress, where they neutralize potentially damaging reactive oxygen species (ROS) (*17–19*). However, they also reduce insect feeding and contribute to mortality of herbivorous insects. Multiple mechanisms for the latter have been suggested, including concentration-dependent deterrence of insect feeding (*20,21*), disruption of Ca^2+^ transport (*22*), inhibition of specific enzymes (*23–25*), and potentially, disruption of the formation of the cross-linked chitin necessary for insect exoskeletons and fungal cell walls.

Carnivorous plants in order Caryophylalles have repurposed the jasmonic acid pathway to also control the prey detection and feeding process (*26*). In the Venus flytrap (*Dionaea muscipula*), wounding and prey capture both induce the same jasmonic acid-regulated response, including trap closure (*27*). Plants have evolved carnivory at least eleven separate times (*28*), suggesting that such co-optation of defensive and other mechanisms to allow prey capture and digestion is relatively straightforward. However, this particular co-option of the ubiquitous jasmonic acid pathway appears to be a distinctive feature of carnivory in Caryphylalles (*29, 30*). Understanding adaptations to plant carnivory and separating the responses to jasmonic acid treatment and digestion of food sources may shed light on other forms of biochemical repurposing in plants and other eukaryotes. In particular, which metabolic changes are a direct response to jasmonic acid or its derivatives, and which require nutrient input from prey?

Our subject organism is the Cape sundew, *Drosera capensis*, which is found in the eponymous Cape region of South Africa. The plant’s elongated leaves bear specialized trichomes tipped with glands that secrete sticky mucilage, which traps prey insects (*31*). Within a few hours of prey capture, the leaf folds and closes around the insect, creating an external stomach into which the plant secretes digestive enzymes to break down its prey into usable nutrients. *D. capensis* is an excellent model organism for studying plant carnivory. It is large and matures quickly, making tissue collection for multiple experiments feasible, and it self-pollinates, meaning that populations maintained in cultivation are relatively genetically homogeneous. It also demonstrates the full range of behaviors characteristic of plant carnivory, including production of specialized structures for prey capture, production of low-pH entrapping mucilage, prey-triggered thigmonasty, production of digestive enzymes, and direct incorporation of nutrients from prey tissues without processing by intermediate species. Past genomic and molecular modeling studies of *D. capensis* have identified numerous families of putative enzymes potentially responsible for these adaptations, but the specific regulatory changes associated with prey capture and feeding remain undercharacterized.

Here, we examine the molecular response of *D. capensis* to prey capture and feeding stimuli, with a focus on the connection between changes in enyzme expression and the production of small molecules associated with signaling, incorporation of prey nutrients, and defense. We exploit the plant’s repurposing of the jasmonic acid pathway to experimentally induce responses in differential expression experiments, allowing feeding behaviors such as leaf curling and transcription of jasmonic acid-regulated genes to be induced without introducing either nutrients or exogenous mRNA from prey. We conduct feeding experiments in which plants are provided with a jasmonic acid signal (a key signal of prey capture) and/or prey nutrients. Using quantitative RNASeq, we examine changes in the expression of proteins associated with a number of feeding-related processes, shedding light on the “molecular machines” mobilized by the plant in response to prey. This is further complemented with mass spectrometry studies of plant metabolites, allowing us to identify small molecules produced during processing of prey tissue (and distinguishing them from molecules produced purely in response to a putative damage or capture signal). Examination of known biochemical pathways associated with observed activities (e.g., production of amino acids and defensive compounds) provides further evidence into the molecular “levers” through which *D. capensis* controls its behavior during the feeding process. In addition to providing insights into plant carnivory, these findings suggest hypotheses for how other, non-carnivorous plants may regulate responses to predation or other environmental stresses.

A brief precis of the remainder of the paper is as follows. We treated unfed *D. capensis* with a solution of jasmonic acid to induce the carnivory response, while monitoring transcriptional regulation of proteins. A transcriptome was assembled, confirming the presence of enzymes predicted from genomic data in previous studies. Differential expression of mRNAs was measured upon treatment with jasmonic acid. Finally, mass spectrometry was used to identify small molecules produced by the plant in the unfed state, after jasmonic acid treatment, and after feeding with either purified protein or insect prey. We find that although jasmonic acid alone stimulates the production of enzymes associated with defensive and digestive processes, many small molecules cannot be produced without input in the form of protein or whole prey.

## Results

### *D. capensis* transcriptome confirms predicted proteins

A short-read genome assembly for *D. capensis* predicted complete matches to 90% (223/248) and partial matches to 99% (245/248) of the core eukaryotic genes in the Core Eukaryotic Genes Mapping Approach (CEGMA) (*32*). In order to facilitate discovering novel enzymes for use in chemical biology and biotechnology applications, molecular models were produced from the genome sequences for particular classes of proteins that are relevant to digestion and remodeling of the leaf tissue, including aspartic proteases (*33, 34*), cysteine proteases (*33, 35*), chitinases (*36*), and esterase/lipases (*37*). As no transcriptome has been previously reported for *D. capensis*, we here provide a genome-guided assembly using RNASeq data obtained from mature leaf tissue (see Methods).

Our transcriptome assembly largely confirms the presence of these putative proteins, including 3/3 of the predicted aspartic proteases, 28/45 cysteine proteases, 6/11 chitinases, and 9/26 esterase lipases. Sequence alignments for selected transcriptome proteins were annotated in detail using a custom tool (*38*); these can be found in the Supplementary Information. Past work on *D. capensis* enzymes focused primarily on molecular modeling and development of hypotheses about the function of these enzymes; the transcriptome provides experimental evidence that the proteins are produced, subject to the caveat that only mature leaf tissue was examined here, meaning that proteins expressed only in other tissues or during earlier developmental phases will not be observed. These estimates are hence conservative. However, the confirmation of 46/85 predicted sequences motivates further investigation into which of these enzymes are involved in the jasmonic acid-regulated defensive/digestive pathway.

### Jasmonic acid treatment induces upregulation of enzymes involved in feeding

We use jasmonic acid to induce the feeding response in mature *D. capensis* leaves, comparing the resulting mRNA expression patterns to those in control leaves treated with distilled water. Jasmonic acid treatment was used to induce the carnivory response in the absence of mRNA from prey, which would complicate the analysis of differentially expressed genes (DEGs). 2170/29939 total genes are upregulated upon jasmonic acid treatment, whereas 2297/29939 are downregulated with an adjusted p-value (p*_adj_*) of less than 0.05. Of those, 815/29939 were at least 2x upregulated in treated samples, while 979/29939 were at least 2x downregulated in treated samples.

### Digestive protease orthologs are upregulated in response to jasmonic acid

Among their other functions, proteases are an important part of plant defensive mechanisms (*39*). Previous work on enzyme discovery from the *D. capensis* genome and *D. muscipula* transcriptome identified several putative aspartic and cysteine proteases in *D. capensis*. Both types of proteases have previously been implicated in digestion in Caryophylalles carnivorous plants (*40–42*) and in plant defenses against biotic stress (*43, 44*). In *D. capensis*, four aspartic proteases are upregulated upon treatment with jasmonic acid (**Figure 1A**). Three of them are nepenthesins, which were first discovered in *Nepenthes* pitcher plants (*46, 47*), where they are upregulated in response to prey capture (*48*). The fourth is related to a known aspartic protease expressed in guard cells (**Supplementary Figure S5**). In *Arabidopsis*, a homolog of this protein is upregulated under drought conditions, where it makes the guard cells more sensitive to the abscisic acid signaling pathway and regulates water loss through the stomata (*49*). In carnivorous plants, abscisic acid signaling has been implicated in the response to wounding, suggesting that there is some cross-talk between the two pathways (*50*).

**Figure 1:**
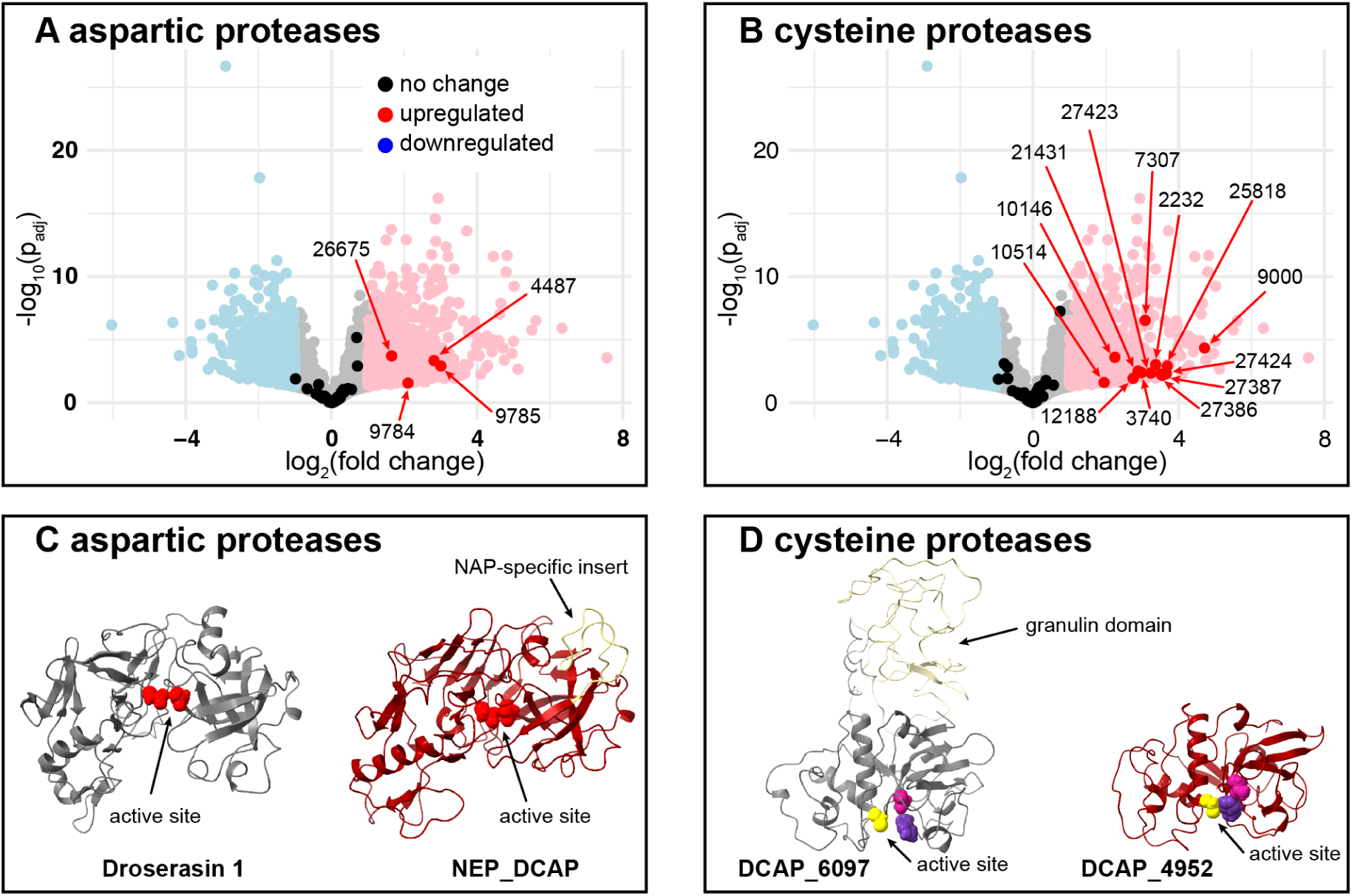
Differential expression of aspartic and cysteine proteases upon treatment with jasmonic acid. (**A**) Jasmonic acid treatment induces upregulation of four aspartic proteases, three nepenthesins (**Supplementary Figure S1**) and one protease associated with drought resistance (**Supplementary Figure S5**). (**B**) Twelve cysteine proteases are upregulated in response to jasmonic acid treatment (**Supplementary Figure S9**). (**C**) Although molecular models show that aspartic proteases share a common fold and active site architecture (*33*) (active site residues shown in bright red), the nepenthesin NEP_DCAP can be distinguished by the presence of the nepthesin-specific insert (NAP, light yellow). The expression level of Droserasin 1 (left), does not significantly change in response to jasmonic acid treatment, which is consistent with its hypothesized role in removal of pro-peptides in vacuoles. NEP_DCAP (right), is one of three nepenthesins upregulated upon treatment with jasmonic acid. (**D**) Both cysteine proteases (models from (*35*)) share the same Cys (yellow) - His (purple) - Asn (magenta) cataytic triad. DCAP_6097 (left) is a cysteine protease with a C-terminal granulin domain (light yellow), a structure typical of proteins involved in degradation of storage proteins during seed germination; it is not upregulated in response to jasmonic acid. DCAP_4952, a Dionain 1 homolog, is upregulated upon jasmonic acid treatment.

In contrast, several other aspartic proteases are present but not upregulated in response to jasmonic acid, including four other nepenthesin paralogs (all sequence alignments are shown in **Supplementary Figures S1 – S8**. For example, Droserasin 1 (model shown in Figure 1B) is homologous to aspartic proteinase A1 in *Arabidopsis thaliana*. This protein is expressed in a variety of tissues, but its primary role is to break down protein inside protein-storage vacuoles in seeds (*51*). Because *D. capensis* has mutiple paralogs of this enzyme (**Supplementary Figure S3**), we had hypothesized that some of them might be related to digestion or protection against pathogens during digestion, and therefore inducible via the jasmonic acid pathway. Instead, differential expression reveals that they are not upregulated upon jasmonic acid treatment and their mRNA level remains constant, consistent with their role in degrading storage proteins during seed germination in other plants (*52*). A molecular model is also shown for the nepenthesin NEP_DCAP, which is upregulated in response to jasmonic acid treatment, consistent with the digestive role of its homologs in *Nepenthes* pitcher plants. It can be identified as a nepenthesin by the presence of a nepthesin-specific insert (NAP), which is shown in light yellow on the model.

Thirteen cysteine proteases are upregulated in response to jasmonic acid treatment (Figure 1C). Sequence alignments are shown in **Supplementary Figure S9**. Overall, these proteases belong to a several diverse lineages with different localization tags, occluding loops, and other sequence and structural features that hint at a wide range of enzymatic activities. However, of the upregulated cysteine proteases, twelve of them are homologs of Dionain 1, a digestive enzyme in the Venus flytrap (*D. muscipula*) (*42, 53*); the thirteenth is more closely related to bromelain from the pineapple (*Ananas comosus*). Models of representative examples are shown in (Figure 1D) DCAP_6097 (left) is a cysteine protease with a C-terminal granulin domain, which indicates its role in degrading storage proteins. As expected based on this function, it is not upregulated in response to jasmonic acid. DCAP_4952, a homolog of Dionain 1, is upregulated upon jasmonic acid treatment. Taken together, these results enable identification of some of the key digestive enzymes in *D. capensis*. Some of the other enzymes in these classes that are not upregulated contain sequence features consistent with roles in cellular protein degradation (*54*), signal sequence removal (*55*), or in specific reproductive functions, such as breakdown of seed storage proteins (*51*), pollen development (*56*), or suppression of programmed cell death during embryo development (*57*).

However, in both enzyme classes, examples can also be found of highly similar paralogs where some are upregulated and others are not, suggesting that selection of digestive enzymes is not solely based on properties of the protein itself. Gene duplication may have led to divergent functionality in different parts of the genome, potentially mediated by different regulatory elements.

### Chitinases are upregulated in response to jasmonic acid; esterases and esterase/lipases show mixed patterns

Jasmonic acid treatment induces upregulation of five chitinases (Figure 2A). Chitinases are an important component of plant defense mechanisms against both insects and fungi (*58*). Treatment with chitosan, the monomeric precursor of chitin, induces production of both defensive enzymes and polyphenolic compounds (*59, 60*). Previous studies of chitinases in *D. capensis* have shown that this plant produces chitinases belonging to both Family 18 and Family 19 (*36, 61*), using the carbohydrate-active enzymes (CAZy) database characterization scheme (*62*). Three Family 18 chitinases (out of seven total found, sequence alignment in **Supplementary Figure S10**) are upregulated in response to jasmonic acid treatment. These enzymes are characterized by a triosephosphatei-somerase (TIM)-barrel fold, with an active site tunnel containing a catalytic glutamic acid that is part of a conserved DXXDXDXE sequence motif (*63*). The chitin polymer is bound by the negatively charged D and E residues as well as several aromatic residues on the opposite side of the tunnel, which enables processive activity (*64*). Family 19 chitinases have much more structural diversity, particularly in the C-rich region and P-rich hinge, both of which are C-terminal to the catalytic domain (*65*). These proteins are often involved in defense against fungal pathogens (*66*). We observe five of these proteins in *D. capensis* (sequence alignment in **Supplementary Figure S11**, two are upregulated in response to jasmonic acid and three are unchanged. Based on genomic data, we predicted a novel two-domain Family 19 chitinase, DCAP_0533, which is present but not upregulated in response to jasmonic acid. A model is shown in Figure 2C **(left)**. Figure 2C **(right)** shows a model of DCAP_4817, one of the single-domain Family 19 chitinases that is upregulated in response to jasmonic acid.

**Figure 2:**
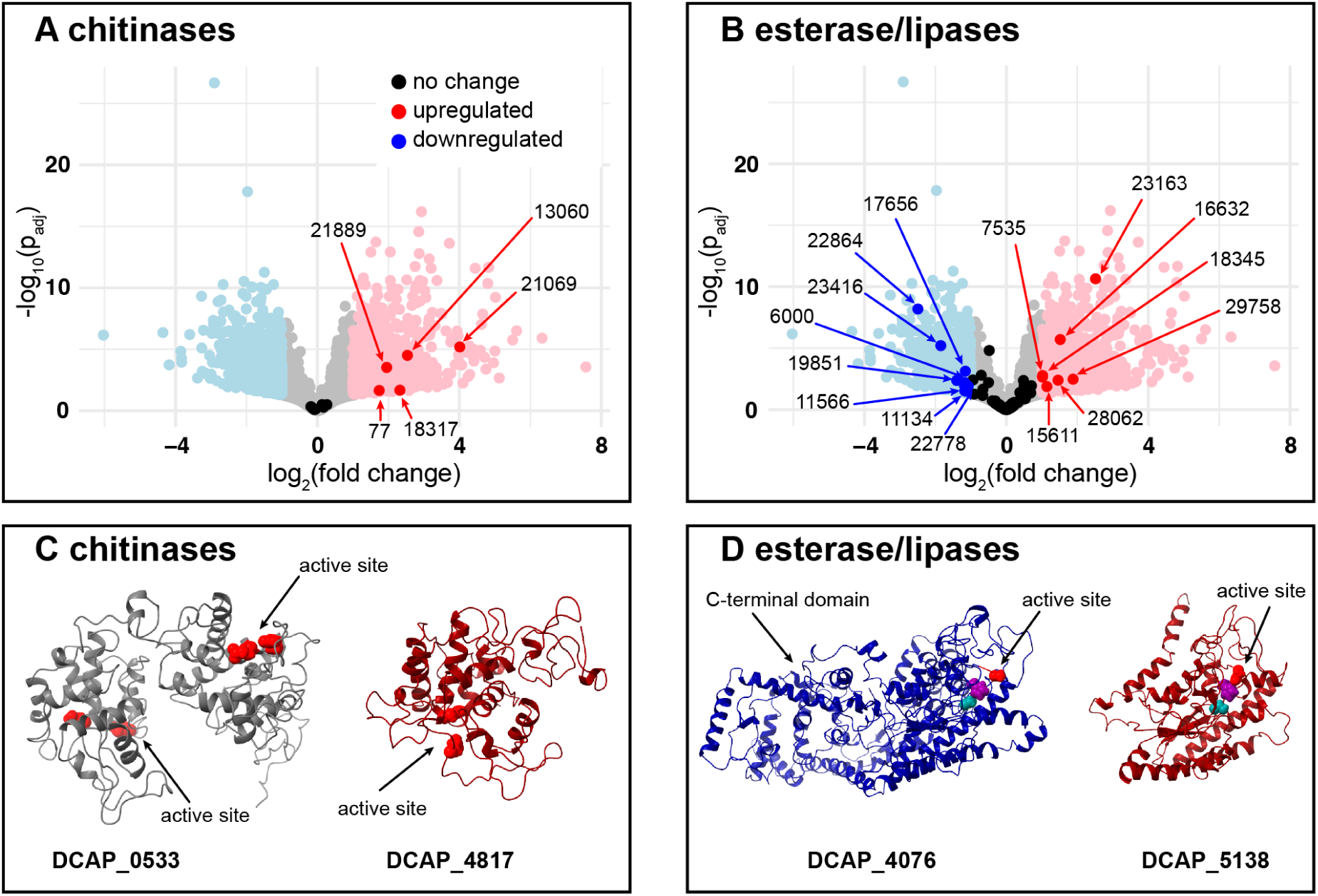
Chitinases and esterases. (**A**) Five chitinases are upregulated upon treatment with jasmonic acid, three from Family 18 and two from Family 19. (**B**) Esterases, a diverse category that includes esterases, acylesterases. and GDSL esterase/lipases, have more complex expression patterns. Seven of them are upregulated and eight are downregulated in response to jasmonic acid treatment.

The expression pattern of esterases is more complex, with seven genes upregulated and eight downregulated in response to jasmonic acid treatment (Figure 2B). Sequence alignments for up- or down-regulated enzymes in this category are shown in **Supplementary Figures S12 – S19**. The pattern where some genes are upregulated and others are downregulated is consistent with the dual roles of esterase/lipases in plants. They can hydrolyze bonds, but they can also run in reverse, synthesizing polyester compounds such as cuticle wax, particularly when embedded in hydrophobic environments (*67*). In the context of prey digestion, it is possible that the plant upregulates esterase/lipases involved in utilizing lipid components from the prey, while downregulating those associated with non-urgent functions such as repair of leaf surfaces, consistent with the observation that jasmonic acid suppresses growth while inducing production of defensive compounds (*68*). In our previous work on GDSL esterase/lipases from *D. capensis*, we defined a measure of active-site flexibility and hypothesized that looser active sites would be correlated with digestive functions, while tighter active sites would correspond to synthesis of polyester compounds (*36*). The limited comparisons available in these data tentatively support that hypothesis, with the upregulated enzymes corresponding to those on the more flexible end of the spectrum. For example, Figure 2D **(left)** shows a model of DCAP_4067, a predicted chitinase with an extra C-terminal domain. It is predicted to have a relatively inflexible active site and is downregulated in response to jasmonic acid treatment. The right panel shows a model of DCAP_5138, which is upregulated upon treatment and is predicted to be more flexible. Although further experimental characterization is needed to fully understand the relationship between active site flexibility and function in these enzymes, the observation of both up-and down-regulated examples in this data set suggests that esterase/lipases play diverse roles in this plant.

### New enzymes were found in the transcriptome

Figure 3A shows the expression levels of neprosins, prolyl endopeptidases first discovered from *Nepenthes* x *ventrata* pitcher fluid (*69*). They have been investigated as an oral treatment for gluten intolerance, as they are active at low pH and reversibly inactivated at pH *>*4.5. The active site has two glutamic acid residues with a bridging water molecule. Despite the mechanistic similarity to aspartic proteases, the overall fold is quite different, with a highly stable *β*-sandwich structure (*70*). We find four neprosin paralogs in *D. capensis* (sequence alignments in **Supplementary Figure S20**. They are all highly similar in sequence; however, two are upregulated upon jasmonic acid treatment while the other two display no difference in expression levels.

**Figure 3:**
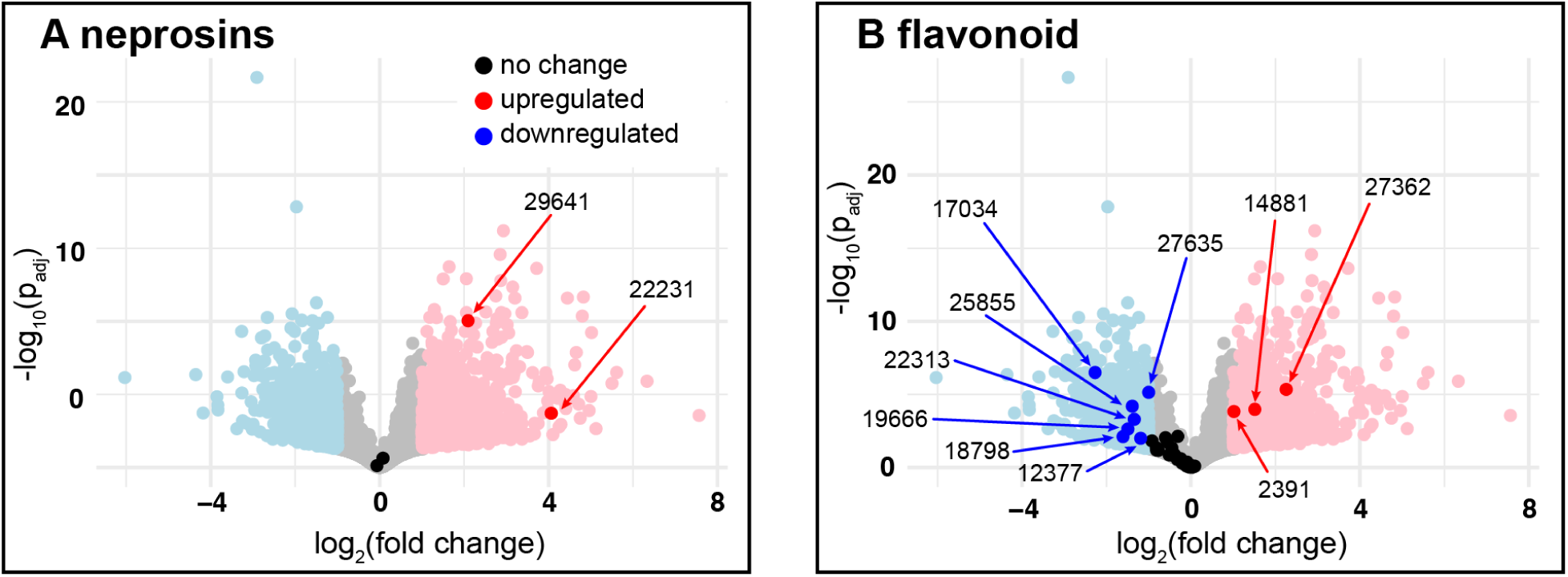
Differential expression of new enzymes upon treatment with jasmonic acid. **A**) Two neprosins are upregulated, while two more show unchanged expression levels. (**B**) Genes whose assigned EC numbers appear in the KEGG flavonoid biosynthesis pathway are largely downregulated.

Figure 3B highlights genes whose predicted EC numbers are present in the flavonoid biosynthesis reference pathway (map00941) in the KEGG pathway database (*71–73*). Flavonoids have many functions in plants, including antioxidant capabilities, protection against UV radiation, attraction of pollinators, and deterrence of herbivores (*74–79*).

Figure 3B shows that most genes that were assigned EC numbers in KEGG’s flavonoid biosynthesis reference pathway were downregulated in response to jasmonic acid treatment. All of the downregulated enzymes are oxidoreductases acting on paired donors (EC 1.14), except for DCAP.17034, which is a caffeoyl-CoA O-methyltransferase. Downregulation of these enzymes is consistent with the idea that a jasmonic acid signal causes the plant to shift resources away from functions like pollinator or prey attraction and UV protection in favor of producing digestive enzymes and growth. Of the 33 genes that were putatively matched with an EC number in the KEGG flavonoid biosynthesis pathway, only 3 (DCAP.2391; EC:1.14.13.-, DCAP.14881; EC:1.14.13.-, and DCAP.27362; EC:2.3.1.133) were at least 2-fold upregulated (FC≥2 and *p_adj_ <*0.05). These are colored red in Figure 3B). Enzymes in class EC:1.14.13 are monooxygenases or hydroxylases: oxidoreductases that incorporate one oxygen atom to the substrate, using NADH or NADPH as the donor. Enzymes in class EC:2.3.1.133 are acyltransferases involved in the phenylpropenoid biosynthesis pathway, specifically shikimate O-hydroxycinnamoyltransferases, although they can also use caffeoyl-CoA, feruloyl-CoA and sinapoyl-CoA, albeit with slower kinetics. Both of these enzyme classes are part of the standard plant toolbox for synthesizing specialized secondary metabolites (*80–82*).

### Nutrient availability and jasmonic acid treatment produce distinct metabolic responses

In order to probe which metabolic responses are triggered by jasmonic acid signaling per se and which are dependent on acquisition of nutrients from prey, we performed a comparative untargeted LC-MS/MS metabolomics experiment, measuring the relative abundances of observable small molecules from *D. capensis* leaves exposed to different conditions (Figures 5**, 6, and 7**). Two different sources of nutrients were provided: protein alone, in this case bovine serum albumin (BSA), and homogenized bloodworms. Bloodworms, which are often sold as fish food, are not worms but the larvae of *Chironomus plumosus*, a small flying insect. Treated leaves were wiped clean and rinsed with deionized water before processing to remove compounds that are on the surface but not absorbed into the leaf tissue. The experimental design is summarized in Figure 5A. There were seven conditions in total: 1) bloodworm homogenate alone, 2) untreated leaf tissue, 3) untreated leaf tissue with bloodworm homogenate added during sample processing under liquid nitrogen, with no time for chemical reactions to occur, 4) leaf tissue incubated with jasmonic acid, 5) leaf tissue incubated with bovine serum albumin 6) leaf tissue incubated with bloodworm homogenate, and 7) leaf tissue incubated with bloodworm homogenate and jasmonic acid. The untreated leaf tissue was collected from the plant without any treatment, providing a resting state baseline. The tissue treated with jasmonic acid provides a condition comparable to that of our differential expression experiments, where the defensive hormone is added to stimulate leaf curling and the production of defensive and digestive enzymes, but without adding nutrients. As an additional control, we added bloodworm homogenate to untreated leaf tissue at the point of grinding the samples in liquid nitrogen with no incubation and no rinsing. This was expected to show approximately the same result as the sum of the untreated leaf and bloodworm samples. The BSA and homogenized bloodworm treatments are intended to test the metabolic response to pure protein and a more complex food source, respectively. A previous timecourse experiment where plants were fed with *Drosophila melanogaster* homogenate showed that although responses begin within six hours of feeding, completion of the digestive process and production of secondary metabolites can last for many days (*83*). Here, we extend these measurements to also investigate the impact of jasmonic acid treatment alone and feeding with a pure protein substrate as well as homogenized insect material.

### Jasmonic acid time-course

The relative abundance of three jasmonic acid derivatives as a function of time after jasmonic acid treatment is shown in Figure 4. Each cell’s color corresponds to the average normalized signal at that time point for that molecule, relative to the most abundant signal in a replicate for that molecule in the entire time course. The bar plot shows the average relative abundance of all three molecules at each time point, and is used to show an average trend. In treating *D. capensis* with JA, we observed a time-dependent increase in three identifiable JA derivatives, namely JA-glucoside, HO-JA-glucoside, and HO-JA-Ile. The glucosides do not appear in the resting state tissue, and we observe a signal maximum at T = 6 hrs. The glucoside derivatives then decrease in signal by 16 hours, although signal remains steady afterward up to 26 hours (the end of the measurement period). We also observe HO-JA-Ile, one of the most common JA derivatives observed in plants, including in a previous study of *Drosera capensis* specifically (*26*). We observed a baseline level HO-JA-Ile even in the untreated plants (T=0 hrs), followed by an increase in abundance between 1 and 6 hours, followed by an approximate 2-fold decrease after 6 hrs back to baseline levels observed in the untreated leaves. These data suggest that treatment of *D. capensis* with jasmonic acid results in a turnover into downstream metabolites on a timescale of hours. Based on these observations and those of Hatcher et al. (*83*), we chose the time point at T=12 hours for a detailed investigation of metabolic responses to jasmonic acid, as this is long enough after the signal induction to allow protein expression and the production of small molecules.

**Figure 4:**
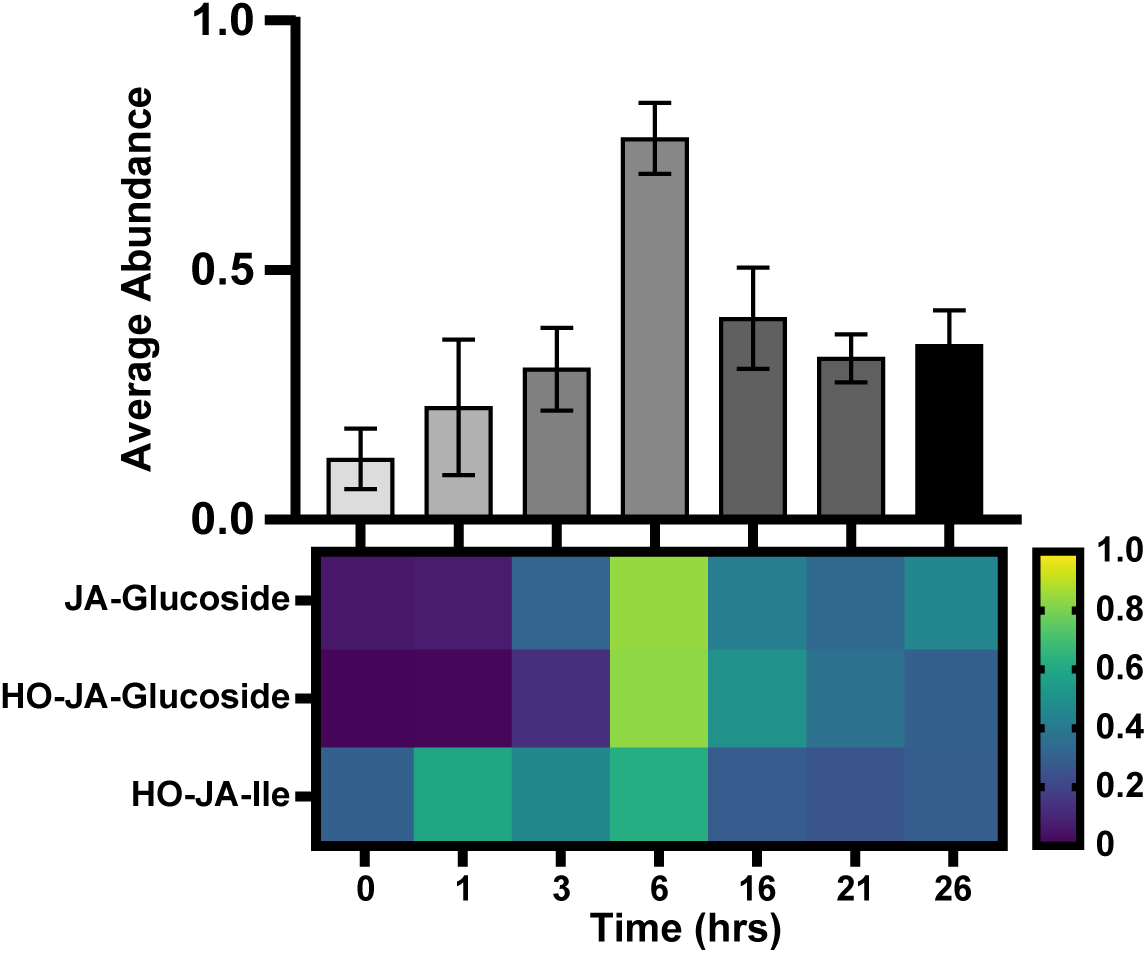
Jasmonic acid derivatives peak at about 6 hours post-treatment. A time-course analysis of jasmonic acid (JA) derivatives identified in *Drosera capensis* (N = 5–9 replicates per timepoint) shows a peak at about 6 hours post treatment.

### Amino acids

Using LC-MS/MS, we observed dramatic changes in the levels of four free amino acids in response to treatment with both food types, but not with jasmonic acid alone (Figures 5B**, Supplementary Figure S21**). A full list of features is tabulated in **Supplementary Data File S1**. Tryptophan, phenylalanine, tyrosine, and arginine were identified with high confidence and significantly changed abundance in response to treatment. All of these amino acids are present in bloodworm homogenate (although the signal for Trp and Phe was low), both alone and mixed with leaf tissue, but not in untreated leaves. The mass spectrometry data presented in Figures 5 – **7** are shown in terms of normalized abundances in order to compensate for the varying ionization efficiencies of different molecules based on their structures and the ionization mode used by the mass spectrometer. For each feature, the abundance value (after correction for sample mass and internal standard signal) is normalized relative to the most abundant molecule in each sample. Therefore, for each cell of the heatmaps, the maximum signal for any one group is almost necessarily higher than the average value by which the cell’s color is determined.

**Figure 5:**
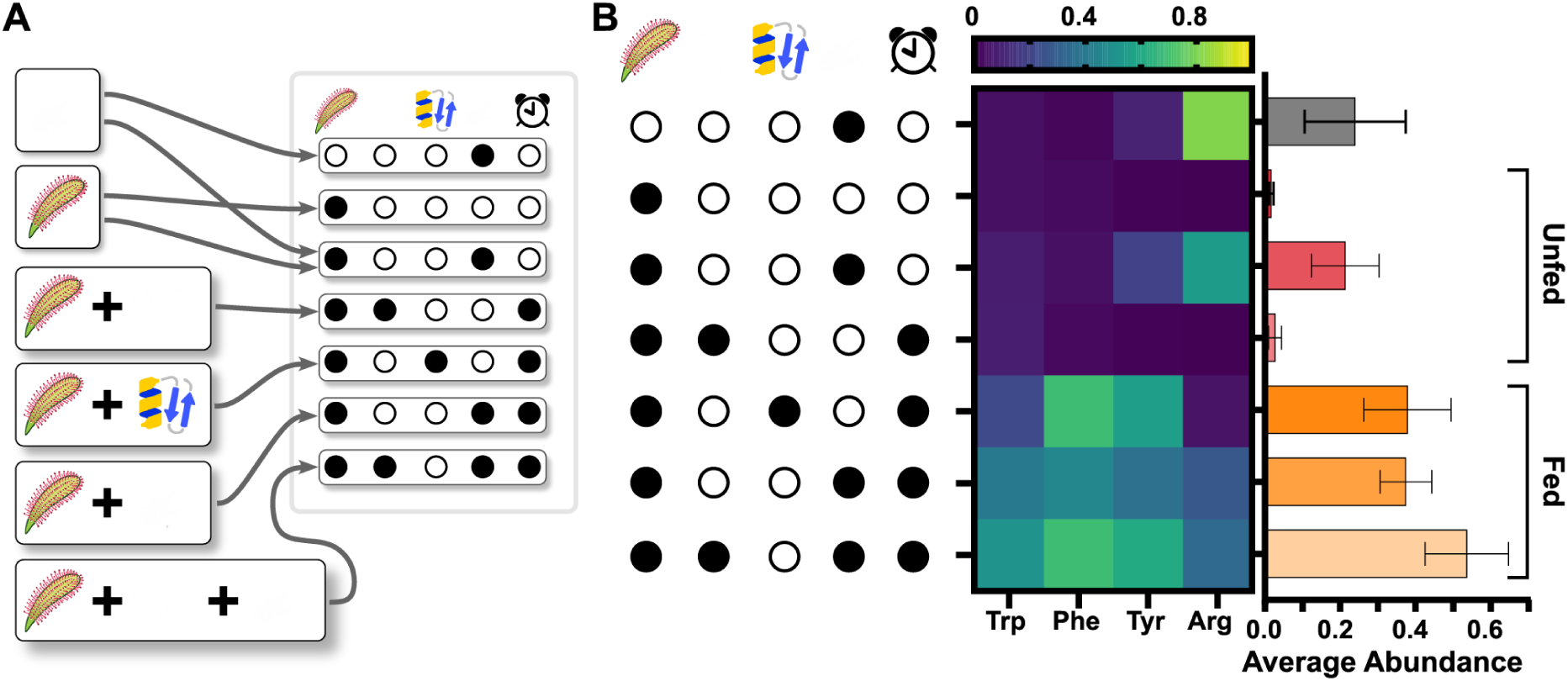
Certain free amino acids increase in abundunce upon feeding. **A**) Experimental design. Leaves were treated with different combinations of jasmonic acid (molecular structure), BSA (protein emoji), and bloodworm homogenate (fly) as indicated by the black and white circles. Time for the leaf to respond to the treatment is indicated by the clock. **B**) Differential treatment analysis of *D. capensis* (N = 7 for each group), tracking the abundance of four amino acids: arginine, tyrosine, phenylalanine, and tryptophan. Each cell of the heatmap shows the average relative abundance of an amino acid, relative to the maximum signal of that amino acid across all replicates. The bar plot shows the average relative abundance of all four amino acids for each treatment. Bars show mean relative abundance, calculated from the relative abundance of all replicates in the treatment group across features, ± 95% confidence interval. Untreated leaves contain very low levels of any of the four free amino acids. Arginine and tyrosine both appear in bloodworm homogenate. All amino acids increased in abundance in response to the consumption of food (either bloodworm homogenate or protein). Arginine and tryptophan levels rose much more in response to bloodworm homogenate, while phenylalanine and tyrosine increased more in response to BSA. When jasmonic acid accompanied treatment with bloodworm homogenate, abundances of all amino acids increased relative to being fed bloodworm homogenate without jasmonic acid.

Free arginine was undetected in the untreated and jasmonic acid treated leaves (at background, 134 counts*min), but showed greatly increased abundance in the leaves incubated with protein treatment (995 counts*min) and bloodworm homogenate (7126 counts*min), resulting in changes of 7.4- and 53-fold relative to the untreated and jasmonic acid treated groups. Arginine displayed the largest relative abundance in the bloodworm homogenate itself, relative to signal in any of the other treatment groups; this indicates that there is at least more arginine present in the complex food than in the resting state leaf tissue, or in leaves that have digested either protein or complex food. There is little difference in arginine abundance between the untreated and protein treated groups, indicating that if BSA is being degraded during digestion as expected, the arginine being freed is either not absorbed or is rapidly converted into other molecules.

Tryptophan increased in abundance in response to all three feeding conditions (bloodworm homogenate, BSA, and bloodworm homogenate + jasmonic acid), although it was also at much lower abundances in the untreated samples and in the bloodworm homogenate. Free tryptophan is not particularly abundant in the undigested food for the plant to absorb directly; this implies that the tryptophan is being liberated using chemical processes. The three fed groups have similar tryptophan abundances; however we do observe a pattern of increasing signal going from protein-fed to bloodworm homogenate-fed to bloodworm homogenate + jasmonic acid. We expect the majority of any observed increase in abundance of an amino acid in this context to come from prey protein digestion by proteases secreted during feeding. Because the masses of protein and bloodworm homogenate received by the plants in these groups were equivalent, the protein-fed group necessarily received a higher total amount of protein *per se*. One might expect a larger increase in abundance of the amino acids when the plant consumes pure protein versus a more complex food source; however, the bloodworm homogenate contains a much larger variety of different proteins. The plant secretes a wide variety of proteases, which are likely to more efficiently cleave a range of individually lower-concentration prey protein sequences than a (relatively) much larger concentration of a single homogenous protein. Further, there may be more efficient absorption of liberated amino acids during complex food digestion based on the presence of other prey molecules, or the plant may have been overloaded with protein, saturating the rate of protease activity.

Tyrosine produced similar results, with background signal of approximately 40 counts*min, but much higher signal in both the protein (2093 counts*min) and bloodworm homogenate (1589 counts*min) conditions, resulting in changes of 52- and 40-fold relative to the untreated and jasmonic acid treated groups. Phenylalanine was present at low but detectable amounts in the untreated and jasmonic acid treated groups, unlike arginine and tyrosine. The untreated and jasmonic acid treated showed an average signal of 91 and 48 counts*min, whereas the protein and bloodworm homogenate treatments showed signals of 3053 and 2075 counts*min, respectively, compared to a background of approximately 14 counts*min. Relative to the untreated leaves, the protein and bloodworm homogenate-treated leaves showed 34- and 23-fold increases in abundance. These patterns suggest that these amino acids, like tryptophan, are being liberated from prey protein. Tyrosine, which is present at a detectable level in the bloodworm homogenate, may also be absorbed directly by the plant. For both phenylalanine and tyrosine, we observe higher abundances in the protein-fed and bloodworm + jasmonic acid-fed groups than the bloodworm homogenate-fed group; this may indicate that abundance is controlled by both the availability of digestible protein and the expression of proteolytic enzymes, which are upregulated by the jasmonic acid treatment.

In addition to free, unmodified amino acids, we also observe apparently deaminated versions of tyrosine (coumaric acid) and tryptophan (3-indoleacrylic acid). Coumaric acid was, like arginine and tyrosine, not detectable in the untreated and jasmonic acid treated groups (background signal 71 counts*min), but showed an increase in abundance in the protein (1194 counts*min) and bloodworm homogenate (947 counts*min) groups, with fold changes of 17- and 13-fold, respectively, relative to the untreated and jasmonic acid-treated groups. 3-indoleacrylic acid was detected above background (7 counts*min) in all samples, but was present in the untreated (226 counts*min) and jasmonic acid-treated groups (211 counts*min) at lower levels than in the protein (464 counts*min) and bloodworm homogenate (1247 counts*min) groups, with changes in the protein and bloodworm homogenate groups of 2- and 6-fold, relative to the untreated group.

### Flavonoids and anthocyanins

The production of flavonoids and anthocyanins under different conditions is shown in Figure 6A. Individual histograms for the selected compounds are shown in **Supplementary Figure S21**, and a full list of the molecules associated with each feature number is provided in **Supplementary Data File S2**. In general, the relative abundances of flavonoids increase in abundance during feeding, although exceptions can be found. Bloodworm homogenate contains effectively no flavonoids, which is expected because these molecules are produced by plants but not animals. Resting state leaf tissue contains a broad variety of these products. In response to treatment with jasmonic acid, *D. capensis* shows a strong overall reduction in abundance of these flavonoids. Given their many defensive roles, this was somewhat unexpected; however their overall increase in abundance when the plant is treated with both jasmonic acid and a food source offers a potential explanation. In the presence of a nutrient source, production of these metabolites is strongly upregulated. Interestingly, the pure protein substrate resulted in an even larger increase in average abundance of flavonoids than with bloodworm homogenate, either alone or with jasmonic acid. This result is consistent with flavonoids being derived from derived from phenylalanine and tryosine via the phenylpropanoid pathway, meaning that pure protein is a more efficient substrate for producing them than an equivalent amount of the more chemically diverse insect homogenate. Annotations for all of the observed flavonoid and anthocyanin molecules are provided in **Supplementary Data File S2**. Three representative molecules are highlighted in (Figure 6B). Their individual histograms showing the response to each condition are shown in **Supplementary Figure S22**. Feature 1070 is putatively identified as quercetin-3-arabinoside. It does not change in abundance in response to jasmonic acid, relative to the resting-state tissue samples, but is upregulated in response to the digestion of protein and bloodworm homogenate, including bloodworm homogenate with jasmonic acid. Feature 807 is putatively identified as myricetin-3-glucose-6’-fucose. This molecule showed near-background levels of abundance in bloodworm homogenate, and a much higher abundance in the resting state plant tissue. The average level decreased slightly in response to jasmonic acid, but with great variability. Its levels increased approximately two-fold in response to the digestion of BSA. A strong, variable increase was also observed in response to the digestion of bloodworm homogenate, both with and without accompanying jasmonic acid treatment. Feature 535 is putatively identified as luteolin-7-glucoside, or cynaroside. Cynaroside was not observed in the bloodworm homogenate, but it was observed in the resting-state plant tissue. Its abundance did not significantly change in response to jasmonic acid treatment; however, its abundance increased approximately two-fold in response to the digestion of any food, with the highest level observed in the protein-fed leaves.

**Figure 6:**
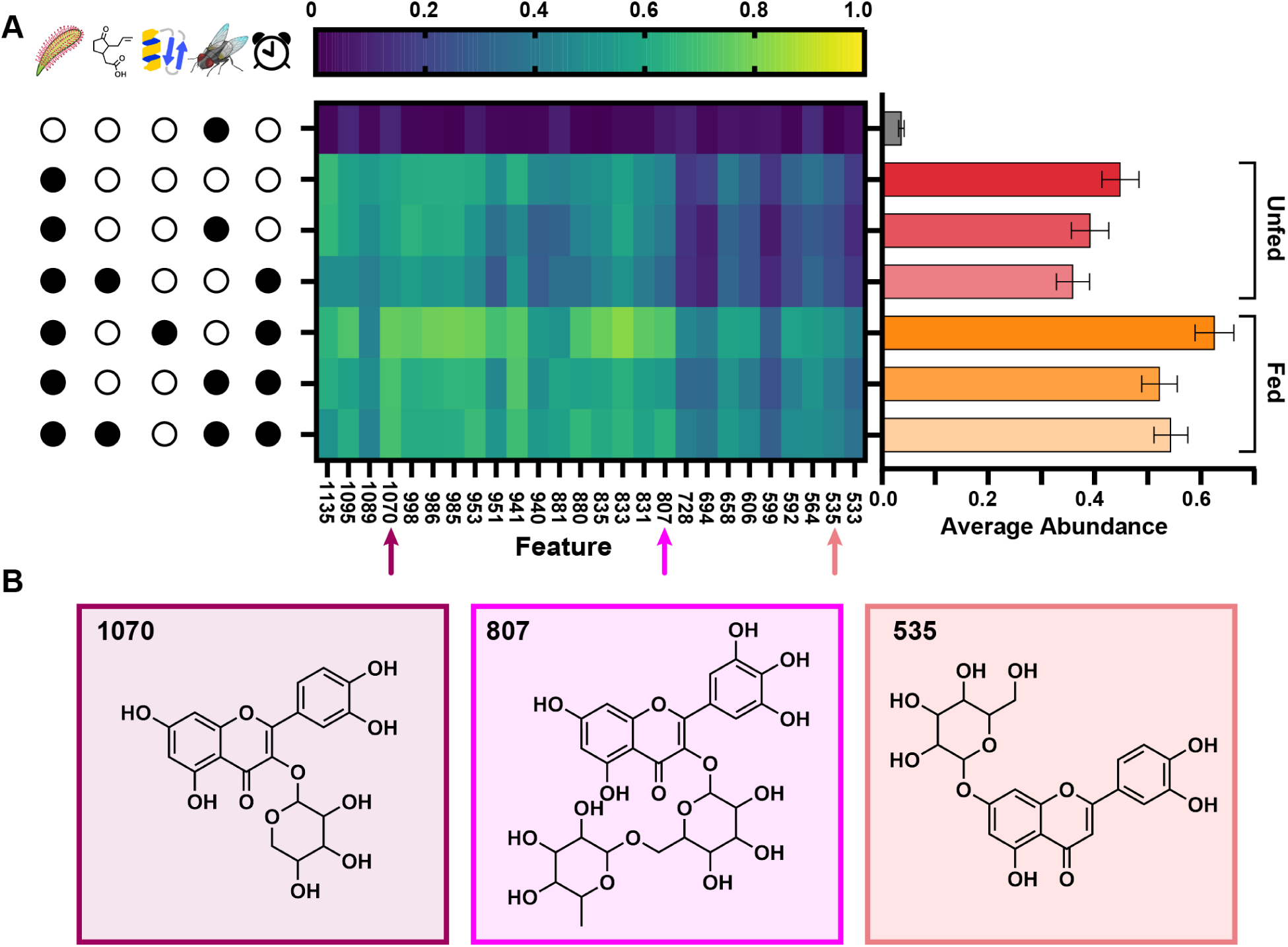
Production of flavonoids and anthocyanins. **A**) A heatmap and bar plot showing the relative abundances of different flavonoids present in D. capensis leaves, in response to various treatments. The color of each cell corresponds to the average replicate abundance (N = 7 for all groups) for *D. capensis* leaves treated with the corresponding substance(s). The bar plot represents the average relative abundance of all flavonoids for a particular treatment. The data show that flavonoids, on average, increase in abundance, when comparing untreated leaves to any treatment where the plant consumes nutrients (bloodworm homogenate or protein). Interestingly, consumption of protein produced a significantly larger increase in the abundance of flavonoids than the consumption of bloodworm homogenate ± jasmonic acid. Treatment with jasmonic acid alone led to a statistically significant decrease in flavonoid abundance, relative to untreated leaves. The three arrows mark example flavonoids chosen as representatives: their individual plots are shown in **Supplementary Figure S22** to highlight their differential abundance in response to various treatments. Bars show mean relative abundance, calculated from the relative abundance of all replicates in the treatment group across features, ± 95% confidence interval. **B**) Structures of three representative flavonoids observed in the dataset.

### Lipids

The production of lipids under different conditions is shown in Figures 7A. Due to space limitations, only selected feature numbers are provided; however, a complete list is given in **Supplementary Data File S3**. Selected molecules are shown in (Figures 7B) Their individual histograms are shown in **Supplementary Figure S23**. Feature 4573 is putatively identified as MGDG(18:2/18:2). This monogalactoglycero-type lipid has two acyl groups (MGDG), which are two identical C18 chains with two double bonds each. These types of lipids are found in cell membranes, but are also effectively hormone precursors. This lipid drops significantly in abundance in response to the consumption of protein, relative to the untreated leaves, but does not drop as much or at all when consuming bloodworm homogenate. Treatment with jasmonic acid alone resulted in little increase in the average abundance relative to the untreated sample, but jasmonic acid + bloodworm homogenate treatment also resulted in an overall drop, to levels about halfway between the untreated and protein treatment groups. The largest effect, produced in response to protein digestion, could suggest that this lipid is tied to proteolysis. The relative response to feeding was the lowest with bloodworm homogenate, followed by jasmonic acid + bloodworm homogenate, and finally protein treatment.

**Figure 7:**
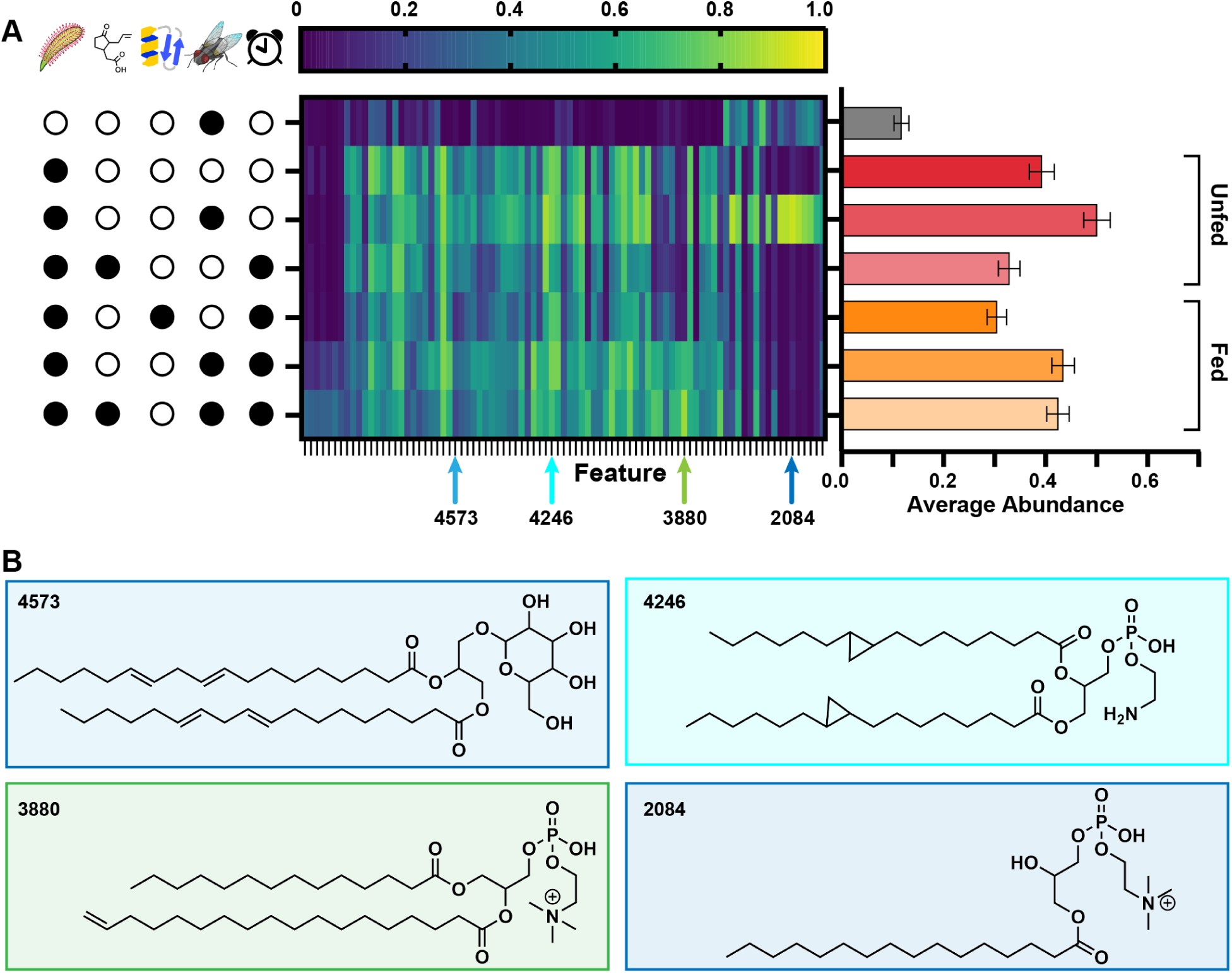
Production of lipids. **A**) A heatmap and bar plot showing the relative abundances of different lipids present in *D. capensis* leaves, in response to various treatments. The color of each cell corresponds to the average replicate abundance (N = 7 for all groups) for leaves that have undergone the corresponding treatment. The bar plot represents the average relative abundance of all lipids for a particular treatment. Only a small fraction of lipids observed in bloodworm homogenate were also found in the plant tissue. We observed a statistically significant decrease in lipid abundance in response to treatment with jasmonic acid, relative to untreated leaves. Interestingly, lipid abundance was reduced even further in response to the consumption of protein, relative to treatment with jasmonic acid. Consumption of bloodworm homogenate, with or without jasmonic acid, resulted in the opposite response – an increase in the abundance of lipids. **B)** Representative lipid structures illustrating different response patterns to the treatments. Individual bar plots showing the abundance for each structure are shown in **Supplementary Figure S23**. Bars show mean relative abundance, calculated from the relative abundance of all replicates in the treatment group across features, ± 95% confidence interval.

Feature 4246 is putatively identified as PE(16:1(cy3)/16:1(cy3)), a phosphatidylethanolamine-type (PE) lipid with two identical C16 acyl groups each bearing a single cyclopropane ring. These types of lipids have been previously found in plants (*84*), where they modulate the biophysical properties of membranes (*85*). PE lipids are generally known to be a large constituent of cell membranes, including associations with membrane proteins. This lipid is only detected in the plant tissue, indicating that it is produced by the plant. Its abundance does not change in response to the consumption of any food type; however, it does decrease in abundance in response to jasmonic acid, with a larger effect in the absence of accompanying bloodworm homogenate. As far as we know, this type of cyclopropane-containing PE lipid has not previously been observed in Caryophylalles; it is previously known from Malvaecaeae, including *Litchi* cultivars (*86*) and cotton *Gossypium hirsutum* (*85*). *Drosera capensis* has two homologs of the cyclopropane fatty acid synthase (**Supplementary Figure S24**), supporting our identification of the lipid. It also has two homologs of diacylglycerol O-acyltransferase 2 (**Supplementary Figure S25**), a critical enzyme for synthesis of triacylglycerol, the major storage lipid in plants (*87, 88*). None of these enzymes is up- or downregulated in response to jasmonic acid treatment (**Supplementary Figures S26, S27**), consistent with the relatively unchanged levels of the lipid itself (**Supplementary Figure S23**).

Feature 3880 is putatively identified as PC(14:0/18:1), a phosphatidylcholine-type (PC) lipid, with one saturated C14 acyl group and a C18 acyl group bearing a single site of unsaturation. The site of unsaturation is drawn at the terminal end of the hydrocarbon chain, but we cannot confidently determine its exact location. This lipid is of low/background abundance in the bloodworm homogenate, and present in low abundance in the resting-state leaf tissue. Its abundance appears to decrease slightly in response to incubation with jasmonic acid alone. Incubation with protein also does not cause a significant shift. However, incubation with bloodworm homogenate, or bloodworm homogenate + jasmonic acid, results in a large upregulation of this lipid. This suggests that this lipid is produced in response to the digestion of non-protein prey nutrients, potentially through the recycling of prey lipids of similar structures. Feature 2084 is putatively identified as lysoPC(16:0), a monoacyl phosphatidylcholine-type (lysoPC) lipid, bearing a single saturated C16 acyl group. This lipid appears to be present in both bloodworm homogenate and resting state leaf tissue, and we observe a striking increase in the abundance of this lipid only in the untreated + bloodworm homogenate control group. Normally, we would expect to see an abundance roughly equal to the combination of the bloodworm homogenate-only and untreated-only group signal, but in this case, it is much higher. This may be indicative of an unknown lipid (such as the di-acyl version of this lipid) being liberated by the combination of the two analyte mixtures during sample preparation.

The relationship between small molecules and enzymes in the general plant flavonoid biosynthesis pathway is summarized in Figure 8, with specific molecules of interest highlighted in the top panel and the full pathway shown in the bottom panel. The expression of enzymes in this KEGG reference pathway (*71–73*) were mostly downregulated (Figure 3B) in response to jasmonic acid treatment. This is consistent with the observation that production of flavonoids and anthocyanins is generally lower in leaves treated with jasmonic acid relative to the untreated control, but feeding with either protein or bloodworm homogenate increases their production (Figures 6A.) This suggests that jasmonic acid is insufficient to trigger upregulation of this pathway: an additional input is needed to increase anthocyanin production. Both EC:1.14.13. (oxidoreductases that incorporate one oxygen atom) and EC:2.3.1.133 (shikimate O-hydroxycinnamoyltransferases) participate in the conversion of p-coumaroyl-CoA (molecule 30) to caffeoyl-CoA (molecule 54) en route to the synthesis of flavonoids like myricetin and luteolin (Figure 8). This conversion of p-coumaroyl-CoA to caffeoyl-CoA is also present in KEGG’s phenylpropanoid biosynthesis pathway (map00940, **Supplementary Figure S28**) (*71–73*). Both p-coumaroyl-coA and caffeoyl-CoA are precursors to versatile phenylpropanoids like methyleugenol, whose uses include pathogen defense and attraction of pollinators (*89*). Notably, the phenolic polymer lignin and its constituent phenylpropanoid monolignols are a structural component of plant cell walls; its biosynthesis is summarized in Tobimatsu *et al.*, 2019 (*90*). Cell wall damage-induced lignin production in *Arabidopsis thaliana* was further shown to be dependent on a balance of reactive oxygen species and jasmonic acid, with jasmonic acid seemingly downregulating lignin biosynthesis (*91*). There are, in general, predicted costs and benefits to the production of secondary metabolites in plants (*92, 93*). Simulations and experiments in apple tree leaves show that concentrations of phenylpropanoids, which include precursors to flavonoid biosynthesis, decrease with increased nitrogen fertilization (*92, 94*). This is consistent with the observation that fast, nitrogen-induced growth is, counterintuitively, associated with a decrease in resistance to pathogen attack (*92, 94*). In the case of anthocyanins, which contain only carbon, oxygen, and hydrogen atoms, the abundance of nitrogen itself cannot be not the limiting factor; more complicated signaling must be at work.

**Figure 8:**
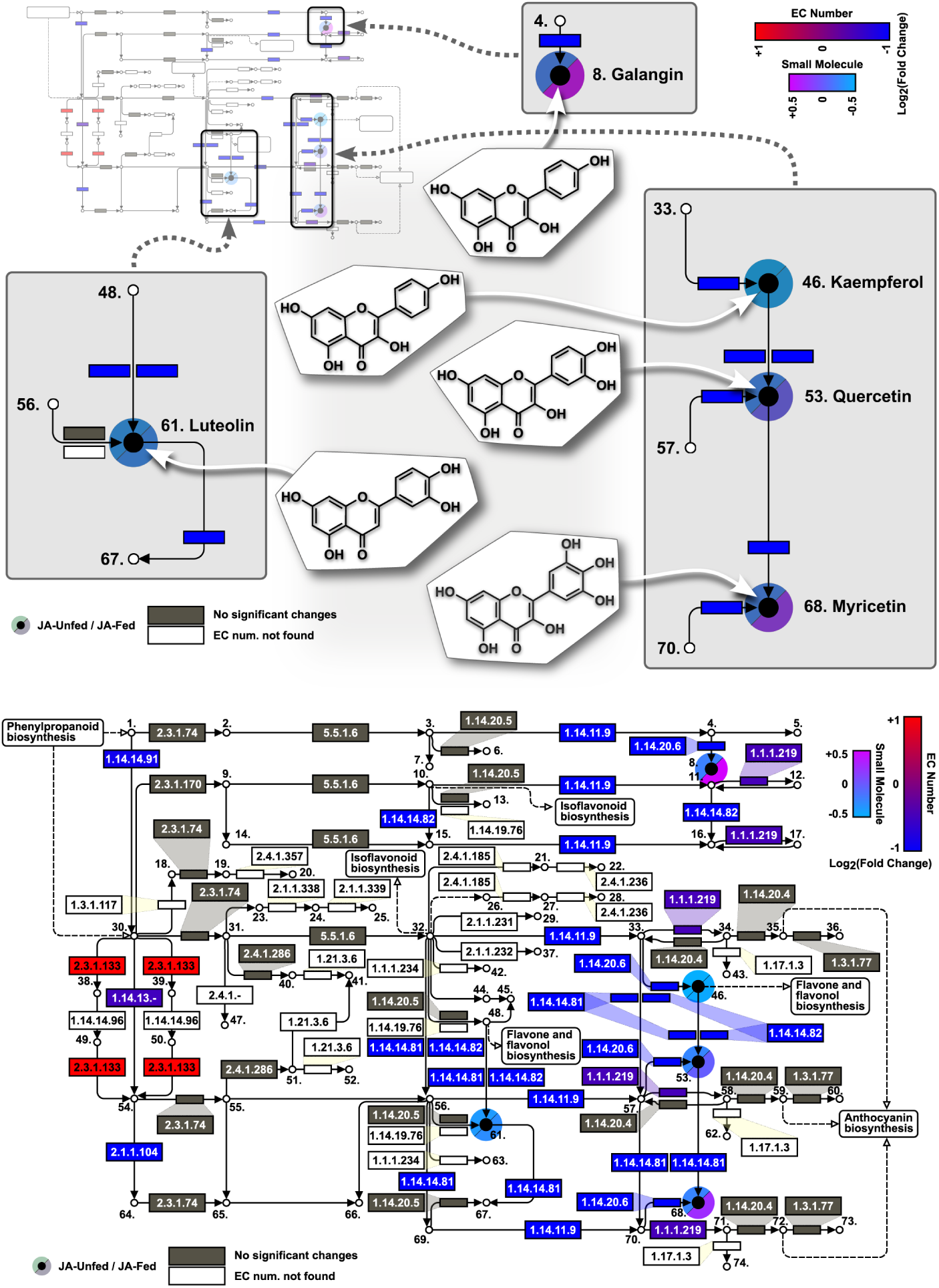
Differential expression of enzymes and small molecules in the KEGG flavonoid biosynthesis pathway (*71–73*) upon treatment with jasmonic acid or jasmonic acid and bloodworm homogenate). bottom: The *log*_2_FC for enzyme expression with jasmonic acid alone are visualized by blue to red colored rectangles. Blue/red coloration represents the average *log*_2_FC of all significant genes assigned the specified EC number. The log_2_FC for small molecule expression with the jasmonic acid alone and jasmonic acid + bloodworm homogenate are visualized in cyan to magenta splitcircles; five small molecules were identified in this pathway: galangin, kaempferol, quercetin, luteolin, and myricetin. EC numbers in grey were found in the transcriptome, but were not assigned to any significant genes. EC numbers in white were not found in the transcriptome. **top:** Structures of the five identified molecules are shown along with close-ups of their positions in the pathway, highlighting the split circles illustrating the differential abundance of each metabolite depending on whether the plant was treated with jasmonic acid alone or with bloodworm homogenate.

## Discussion

The roles and production of flavonoids in carnivorous plants mirror those in their non-carnivorous counterparts. In addition to pollinator attraction and ties to nitrogen deficiency, it is hypothesized that trap coloration in both visible and UV spectra may contribute to prey attraction in carnivorous plants, although many invertebrates do not see red shades as distinct from green. Evidence for and against this hypothesis is discussed in Hatcher *et al.*, 2019 (*95*). There is a repeatedly demonstrated link between nitrogen deficiency and flavonoid accumulation (*74, 96, 97*), which is potentially relevant to the evolution of plant carnivory in nitrogen-limited environments, where the benefits of additional nutrients outweigh the cost of maintaining the molecular machinery necessary for prey capture and digestion. In *D. capensis* specifically, the levels of two flavonoids, quercetin and myricetin, were shown to increase when grown under nitrogen-poor conditions (*98*).

Secondary metabolites have previously been shown to be produced in response to prey capture in *Drosera capensis* (*83*). Here we distinguish between upregulation of enzymes in response to a jasmonic acid signal and the metabolites produced in response to nutrient inputs from prey. As in the previous studies, we observe an increase in aromatic amino acids in response to the consumption of bloodworm homogenate or protein, but not in response to jasmonic acid alone. We also observe broad upregulation of flavonoid biosynthesis in response to feeding but not jasmonic acid alone. Some of the molecules observed by (*83*) were also found in our dataset but were attributed to the bloodworm homogenate itself, rather than a metabolite produced by the plant during digestion (e.g., carnitine). Comparing to the results summarized in the review (*99*), we observe many of the phenolic acids shown to be present across different *Drosera* subspecies. These include monogalloyl glucoside, trigalloyl glucoside, and di-O-methylellagic acid.

Even genes that did not pass FC thresholds for coloration in Figure 3 (see **Materials and Methods**) had negative *log*_2_FC values. This suggests that jasmonic treatment leads to a general reduction in flavonoid biosynthesis, an observation that is corroborated by a reduction in kaempferol, quercetin, myricetin, luteolin, and galangin for leaves treated with jasmonic acid in the absence of nutrients (upper left half of transparent circles in Figure 8).

In general, our data show that phenylalanine, tyrosine and other amino acids are upregulated when substrate (BSA or bloodworm homogenate) is provided (**Supplementary Figures S21** and **S28**); enzymes that break down phenylalanine and tyrosine (phenylalanine ammonia lyase - PAL, EC:4.3.1.24 - and phenylaline/tyrosine ammonia lyase - PTAL, EC:4.3.1.25) are downregulated in response to jasmonic acid treatment (**Supplementary Figure S28**); enzymes involved in flavonoid biosynthesis are mostly downregulated in response to jasmonic acid treatment (Figure 8); enzymes involved in other phenylpropanoid biosynthesis, including structural molecules like monolignols, are more upregulated than are those along the flavonoid biosynthesis pathway (Figure 8 and **Supplementary Figure S28**); and flavonoid production is generally downregulated in response to jasmonic acid treatment alone, though many flavonoids are upregulated when bloodworm homogenate is added. We therefore conclude that when *D. capensis* receives a jasmonic acid signal, it upregulates the production of digestive enzymes while downregulating the production of most secondary metabolites. When these changes in enzyme regulation are followed by an influx of nitrogen and amino acids (represented by the small molecule reaction to jasmonic acid plus bloodworm homogenate), the plant produces a variety of growth-related and defensive secondary metabolites.

## Conclusion

*Drosera capensis* was shown to have a differential response to treatment with jasmonic acid as compared to food, either pure protein or more complex insect tissue. *D. capensis* leaves displays an increased abundance of arginine, phenylalanine, tyrosine, and tryptophan in response to the consumption of bloodworm homogenate or protein (in the form of bovine serum albumin). Flavonoid abundance increases in response to feeding, but decreases in response to treatment with jasmonic acid alone. The reduction of flavonoid abundance in response to jasmonate treatment was rescued when the plant was given bloodworm homogenate alongside the food, suggesting further that jasmonic acid plays a key role in the upregulation of carnivory-related pathways, but can cause downregulation of these same pathway products when nutrient-containing substrates (food) are lacking when the plant recieves jasmonic acid. We also observe broad upregulation of lipids in *D. capensis* in response to the consumption of bloodworm homogenate, however feeding with protein or treatment with jasmonic acid result in downregulation of lipids. This suggests that lipid abundance overall only increases when the plant is digesting food that is more easily converted to lipids, perhaps for growth of energy storage.

## Acknowledgments

The authors thank Felix Grün and Benjamin Katz for excellent management of the UCI mass spectrometry facility, and Melanie Oakes and the UCI High-Throughput Genomics Center for assistance with collecting RNASeq data. This work utilized resources of the UCI Genomics Research and Technology Hub (GRT Hub) parts of which are supported by NIH grants to the Comprehensive Cancer Center (P30CA-062203) and the UCI Skin Biology Resource Based Center (P30AR075047) at the University of California, Irvine, as well as to the GRT Hub for instrumentation (1S10OD010794-01and 1S10OD021718-01). We thank Veronika César and Ethan Kocak for providing the leaf and fly diagrams, respectively.

## Author contributions

Z.G.L. participated in study design, collected and analyzed mass spectrometry data, acquired funding, and wrote the manuscript. G.R.T. participated in study design, collected and analyzed transcriptome and differential expression data and wrote the manuscript. R.W.M. and C.T.B. participated in study design, acquired funding, analyzed data, and wrote the manuscript.

## Data and materials availability

The raw RNA data are available in GenBank under BioProject PRJNA1101963. Differential expression and metabolomics data can be found in an Open Science Framework (OSF) repository: https://osf.io/h6beu.

## Supplementary Materials for

### Materials and Methods

#### Plant cultivation

Seed-grown *Drosera capensis* specimens were grown under laboratory conditions, watered with reverse osmosis (RO) water, fed with freeze-dried bloodworms, and kept under plant lamps on a 12-hour light/dark cycle.

#### Metabolomics

Bloodworm homogenate (BWH) was prepared by grinding 194.5 mg of freeze-dried bloodworms into a fine powder using a mortar and pestle, followed by the addition of 1945 *µ*L of nanopure water and additional grinding. Treatment of leaves using BWH involved placing three 10 *µ*L droplets spaced evenly along the leaf blade, followed by the spreading of the homogenate across the leaf blade using the pipette tip, while also disturbing the trichomes of the leaf blade. This resulted in the deposition of 3 mg of bloodworm mass to each leaf.

Jasmonic acid (JA) treatment was carried out similarly to the BWH treatment above, but used 500 *µ*M jasmonic acid in nanopure water, in place of BWH. Protein treatment was also carried out similarly to BWH and JA treatments, but used 100 mg/mL bovine serum albumin in water instead of BWH or JA. The JA + BWH treatment used bloodworm homogenate prepared with 500 *µ*M JA as the solvent instead of nanopure water.

Extraction solvent was prepared by making 90% MeOH/10% nanopure water (v/v), containing 50 uM of each N-lauroylsarcosine and 3-chloro 4-methoxyaniline as internal standards. Dilution solvent was prepared by making 0.1% formic acid in water (v/v) containing 5 uM of each chloramphenicol, 8-chlorotheophylline, ampicillin, carbenicillin, and spectinomycin, as well as 250 nM of each: PEG200, PEG400, PEGMME750, PPGB2APE400, and PPGB2APE2000. The standards in the dilution solvent were used mainly for retention time alignment between samples, while the N-lauroylsarcosine in the extraction solvent was used for signal normalization between samples.

*D. capensis* leaves used for untargeted metabolomics experiments were treated in six different groups: 1) Untreated, 2) Untreated + BWH, 3) JA Treatment, 4) Protein Treatment, 5) BWH treatment, and 6) JA + BWH Treatment. A seventh group containing no plant tissue, BWH Only, was also prepared.

The “Untreated” leaf group comprised leaves collected from resting-state plants which were not treated with any substance prior to collection. The “Untreated + BWH” group was also comprised of untreated leaves, but BWH (30 *µ*L) was added directly to the extraction solvent during sample homogenization after leaf collection, with the intent of showing which molecules were present in the BWH without giving the plant the opportunity to act on the substrate. This is paired with the “BWH Only” group, which contains no leaf tissue - BWH (30 uL) was added directly to the same volume of extraction solvent (2 mL) as the leaf samples; theoretically, the “BWH Only” and “Untreated + BWH” group should show the same signal intensities for molecules which only appear in the BWH, if the mixing of leaf homogenate in extraction solvent with BWH results in no metabolism, as desired. The “JA Treatment”, “Protein Treatment”, “BWH Treatment”, and “JA + BWH Treatment” all involved the treatment of leaves with their corresponding solutions 12 hours prior to collection.

After the appropriate designated incubation time, leaves were collected from the plants using clean stainless steel tweezers and scissors to cut the leaf blades from the plant with minimal plant stalk attached. The leaves were then wiped using KimWipes to remove the bulk of the mucilage and treatment residue, and then washed in batches in a 50 mL plastic conical vial. The leaves were washed five times using about 40 mL of RO water, shaking the tube for approximately 15 seconds each time, to remove the remaining mucilage and any water-soluble surface contaminants. The leaves were then patted dry using paper towels and KimWipes, before being placed in individual labeled 1.5 mL microcentrifuge tubes in order to obtain the mass of each leaf for normalization purposes. The average leaf mass across all replicates was 47.9 ± 3.1 mg (95% CI, N = 42), with a median of 47.8 mg.

After the mass of each leaf was found and noted, the leaves were individually ground in a mortar and pestle after flash freezing in liquid nitrogen, until a fine paste formed (about 1 minute with vigorous homogenization). Extraction solvent (2000 *µ*L) was added and mixed in to the slurry with further vigorous mixing in the mortar and pestle. The thin slurry was then pipetted into microcentrifuge tubes, and all samples were centrifuged at 14.1k g for 2 minutes. Supernatant was then diluted ten-fold (100 *µ*L + 900 *µ*L) with dilution solvent. The samples were then mixed vigorously by vortexing, followed by additional centrifugation at 14.1k g for 90 seconds. Supernatant was then transferred to glass pre-slit screw-top LC-MS vials for analysis.

For each sample group (Untreated, JA Treatment, etc.), a pooled sample of all replicates in that group was made by combining equal parts of each replicate’s methanolic extract together, effectively forming an average sample for each group. These grouped samples were then used to make an ‘all-group’ mixed sample, by again combining equal parts of each group’s average sample together, again prior to mixing with dilution solvent. The ‘all-group’ extract sample was then used to make a dilution series, formed by serially diluting the methanolic extract 4-fold with extraction solvent (containing the two internal standards, namely N-lauroylsarcosine), keeping the internal standard concentration the same but diluting all other components by 4, 16, 64, 256, and 1024-fold.

All methanolic extracts were stored at −80 ^◦^C until the day of analysis, when they were diluted with dilution solvent and further processed and loaded into vials as explained above.

All samples were run using matching chromatographic methods. Samples were applied to an Acquity Premier CSH Phenyl-Hexyl 1.7 *µ*m VanGuard FI column (2.1×50 mm), with a 30 minute gradient of 0.1% FA (A) and acetonitrile (ACN with 0.1% FA; B). The gradient was constructed as follows: 0-2 min, 0% B; 2-25 minutes, linear ramp to 100% B; 25-27 minutes, isocratic 100% B; 27-28 minutes, linear ramp down to 95% B; 28-29 minutes, linear ramp down to 0% B; 29-30 minutes, isocratic 0% B. The flow rate was 0.400 mL/min across the entire gradient, except at 30 minutes when it was reduced to 0.050 mL/min.

All data were acquired in centroid-mode, with a LockMass correction applied. LockSpray (LeuEnkephalin, 556.2771 m/z for positive mode; peak detection set to target +/- 0.25 Da) was configured to spray at 10 uL/min during the gradient on a separate channel, collecting a 0.25 second scan every 10 seconds, with the LockMass correction being applied using an average of the last three LockMass scans.

Samples associated with the feeding experiment were analyzed in positive mode. Fast Data Directed Acquisition (FastDDA or FDDA) experiments were used to collect both MS1 and MS2 data on all samples. The FDDA experiments were set up to collect a cycle of scans: one MS1 ‘parent’ scan, to identify molecules of interest, followed by a single MS2 scan on the most abundant singly-charged precursor of interest as identified in the MS1 scan - this cycle was repeated over the full chromatography period. The experiments here consisted of a one-second cycle of a single 0.2 second MS1 scan across the full mass range of interest (50-2000 Da), followed by one MS2 scan for 0.8 seconds. One replicate from each of the seven groups was run in sequence with one other replicate from each group, and in each block of replicates the order of the groups was shifted by one; meaning, the first block ran A-B-C…-G, the second block ran G-A-B…F. Each block utilized a new exclusion list, which was expanded at the end of each sample based on what features were sent for fragmentation. This was done using “AutoCat”, a script provided by Waters (the manufacturer of the Xevo) to allow iterative expansion of exclusion lists from run-to-run, in order to maximize diverse sampling of precursors while minimizing oversampling of the same precursor from sample-to-sample. A collision energy ramp of 10-100V was applied during all MS2 acquisitions.

Waters RAW files were converted to mzML files using the MSConvert application from ProteoWizard. In order to remove LockMass scans prior to processing in MZMine, a subset filter was applied as necessary. For FDDA experiments, with MS1 (1) and MS2 (1) scans in every cycle, the subset filter was set for MS level = 1-2, and scan events 1-2 (excluding the LockMass channel/event 3). Otherwise, default settings were used. Resulting mzML files were renamed to simplify data processing.

The mzML files were loaded into MZMine 4 for analysis, and analyzed using a batch format. The batch file used for data processing is included, as are screenshots of each of the settings pages. The end result of data processing was the production of a few files: 1) a .csv file containing quantitative area values for each feature in each sample (from the Metaboanalyst export), 2) a .mgf file for structural determination of features using SIRIUS.

The Metaboanalyst export .csv file was briefly processed in Excel, where a single row was added below the sample name row and labeled “Tissue Mass (mg)”, and each sample had its previously determined tissue mass noted. For grouped samples, the average tissue mass for the relevant group was used. For samples which were prepared using a mix of grouped samples, the average of the average mass was used. The “BWH Only” group, containing no leaf tissue, used the average mass of all groups, since this was the best way to keep the normalization reasonable. The feature ID for N-lauroylsarcosine was identified and noted, for use in the R script processing.

The .mgf output from MZMine was loaded into SIRIUS 6 for putative structural determination of all molecules in the datasets for which MS2 information was collected. SIRIUS 6 was used with the settings shown in Figure X. SIRIUS was run locally on personal hardware. The SIRIUS project file was saved locally and accessed via the API using R. Features of interest, as far as ‘reasonable structural identifications’, were limited to features with CSI FingerID scores (usually) between 0 and −100, and Tanimoto similarity scores of ≥ 70%.

Once the datasets had been processed in MZMine and SIRIUS, the .csv file and SIRIUS project were utilized by the accompanying R script to perform multiple functions, described briefly as: 1) import the quantitation data from the .csv file exported by MZMine, 2) calculate and apply a signal intensity correction to each sample based on the signal of the internal standard (N-lauroylsarcosine), 3) calculate relevant statistics such as feature mean, standard deviation, etc. by sample group, 4) access the SIRIUS project using API calls to import relevant structural information, 5) filter structural hits by CSI Finger ID score and Tanimoto similarity, 6) produce summary tables of both identified and unidentified features in the dataset, 7) determine statistical significance of both identified and unidentified features, and 8) produce and export basic plots of statistically significant features, with structures shown where features had reasonable structure hits.

The final quantitation table “MetaboanalystUploadExport” contains a full list of features, all normalized to the internal standard signal and corrected based on the tissue mass in each sample. The dilution series of quality control samples was used to gap-fill otherwise empty cells, based on the minimum signal intensity detected for each feature in a row (not across all features), as a more accurate and realistic means of estimating the limit of detection for unknown compounds, as compared to arbitrarily gap-filling empty cells with 1s or filling in cells with some fraction of the lowest detected signal for a feature across samples.

The JA treatment time-course experiment was carried out similarly to the 12-hour differential treatment experiment, but leaves treated with JA were left to incubate for an appropriate number of hours prior to collection and sample processing. Samples were analyzed using a downscaled 15-minute gradient on an identical column and instrument, with similar experimental conditions. Tune file settings for negative mode were identical to positive mode, except the capillary voltage was lowered to 2 kV. Negative mode was used for these samples due to greater ionization efficiency of JA and derivatives in negative mode.

### RNA Extraction and Sequencing

#### Experiments

Leaves were treated with either: three 10*µ*L droplets of water (control) or three 10*µ*L droplets of 500 *µ*M jasmonic acid (Sigma-Aldrich, J2500) (treated). Leaves treated with jasmonic acid exhibited digestive behavior, but not those treated with a water control; for the purposes of this study, “digestive behavior” is defined as a curling of both the trichomes and the leaf itself several hours after treatment.

#### Extractions

After 10–15 hours, leaves were harvested and RNA was extracted using a modified version of the Spectrum Plant Total RNA Kit protocol A (Sigma-Aldrich, STRN10) (described in Kalinowska et al., 2012 (*100*)). All mortars, pestles, pipette tips, metal spatulas, and metal scissors were treated with RNaseZAPTM (Sigma-Aldrich, R2020), rinsed with nuclease-free water (DEPC-treated), and autoclaved prior to use.

#### Sequencing

Quality control, library preparation, and sequencing were performed by the UCI Genomics High Throughput Facility (https://genomics.uci.edu/). Six control and six treated samples were chosen based on RNA integrity number (RIN) and total amount of RNA, as determined by an Agilent bioanalyzer (control RINs: 8.9, 9.5, 9.5, 8.9, 8.7, 8.9; treated RINs: 9.2, 8.6, 9, 8.8, 9, 8.8). Libraries were prepared using the Illumina TruSeq mRNA stranded library construction with unique dual indexes. Paired-end reads of length 150 bases were retrieved using an Illumina NovaSeq 6000 (control: 20,425,830 paired-sequences, 18,548,472 paired-sequences, 17,503,172 paired-sequences, 22,634,376 paired-sequences, 21,894,571 paired-sequences, 29,847,554 paired-sequences; treated: 21,658,099 paired-sequences, 19,175,304 paired-sequences, 17,379,155 paired-sequences, 23,587,565 paired-sequences, 22,451,990 paired-sequences, 22,413,643 paired-sequences).

### Transcriptome Assembly

The raw data for this assembly can be found under BioProject PRJNA1101963.

#### Initial assembly

The existing *Drosera capensis* draft genome (GenBank: GCA_001925005.1 (*33*)) was first indexed using STAR 2.7.10a (*101*), with genomeSAindexNbases 12. Each set of reads was then aligned individually to the indexed genome (using STAR 2.7.10a, with –twopassMode Basic, –outFilterMismatchNoverLmax 0.1, –outFilterMismatchNmax 30 (*101*)); twelve BAM files were outputted, sorted by coordinate. XS tags, important for stranded data, were added to the resulting BAM files using Samtools 1.15.1 (*102, 103*). These tagged BAM files were then individually assembled using StringTie 2.2.1 (*104*); the twelve resulting GTF files were merged using StringTie 2.2.1 (*104*) to produce an initial transcriptome assembly (GTF format). A FASTA-formatted version of this transcriptome was also generated using gffread from the Cufflinks 2.2.1 suite (*105*).

#### rRNA removal

Transcripts from this StringTie assembly were queried against a custom BLAST v2.12.0 (*106*) database of twelve genomic, chloroplast, and mitochondrial rRNA sequences from various plants (NCBI (*107*) organism/taxid/description/Gene ID: Solanum lycopersicum (tomato)/4081/4.5S ribosomal RNA/3950433, Solanum lycopersicum (tomato)/4081/5S ribosomal RNA/3950435, Solanum lycopersicum (tomato)/4081/16S ribosomal RNA/3950430, Solanum lycopersicum (tomato)/4081/23S ribosomal RNA/3950431, Solanum lycopersicum (tomato)/4081/5S ribosomal RNA/34678306, Suaeda glauca/397272/12S ribosomal RNA/70597357, Cannabis sativa/3483/large subunit ribosomal/27215487, Solanum lycopersicum (tomato)/4081/5S ribosomal RNA/112940756, Solanum lycopersicum (tomato)/4081/5.8S ribosomal RNA112940380, Arabidopsis thaliana (thale cress)/3702/18S ribosomal RNA/3767991, Solanum lycopersicum (tomato)/4081/25S ribosomal RNA/108175346, Solanum lycopersicum (tomato)/4081/28S ribosomal RNA/112940383). All transcripts that matched any of the twelve subject rRNA sequences with an e-value less than 1e-5 were removed from the assembly. Remaining transcripts with greater than 10% N content, greater than 14 Ns in a row, or fewer than 200 bp were also removed. Further, nineteen transcripts were removed using NCBI’s transcriptome shotgun assembly (TSA) contamination screening process. This filtered version of the assembly was used for the analyses in this paper. A separate version of the assembly was submitted to NCBI’s TSA database, in which leading and lagging Ns were cut from transcripts (GKUK00000000).

This assembly contains 29939 genes and has a BUSCO v5.7.1 (*108–110*) score of 87.3% (complete BUSCOs), using the embryophyta dataset (embryophyte_odb10), which contains 1614 BUSCOs.

#### Gene counts

Gene counts for each sample were acquired by re-aligning raw reads to the indexed genome, using the new GTF-formatted transcriptome as an additional reference; the same settings as in the initial alignment were used along with –quantMode GeneCounts. Read counts in the resulting *ReadsPerGene.out.tab files were used in differential expression analyses. The number of uniquely mapped reads after this step were: control: 12359436, 15139840, 13388650, 13099403, 16157201, 21598995; treated: 16289554, 14538701, 13383070, 17713204, 16426194, 17268297.

### Differential Expression

Differential expression analysis was performed using the DESeq2 v1.46.0 package (*111*) in R v4.4.2 (*112*). Raw reads from the *ReadsPerGene.out.tab files were combined into a single table and passed to DESeq2 in R to generate a DESeqDataSet object. At this time, genes whose counts across all twelve samples totaled less than 100 were removed, leaving 24980 genes. A results object was generated with an alpha cutoff of 0.05, and 157 genes were identified as outliers via Cook’s distances (*113*); these genes were left in the results object, but their p-value and adjusted p-value were set to NA. The default DESeq2 Cook’s distance cutoff was used: the .99 quantile of the F(p, m-p) distribution, where p and m are the number of coefficients and the number of samples (*111, 112*), respectively. The log_2_ fold change (*log*_2_FC), here defined as the *log*_2_ of treated expression (JA-treatment) over control expression (water-treatment), of each gene was shrunk using the apeglm function (*114*). These results were saved and used in downstream analyses.

In this set, there were 2170 upregulated (positive *log*_2_FC), and 2297 downregulated (negative log_2_FC) genes with an adjusted p-value (padj) of less than 0.05. Of those, 979 had a *log*_2_FC less than or equal to −1, and 815 had a log_2_FC greater than or equal to +1; these correspond with a fold change of at least 2x in either direction. Volcano plots were generated using ggplot2 v3.5.1 (*115*) in R v4.4.2 (*112*); all volcano plots included all genes except the 157 outliers identified with Cook’s distances. Points were colored based on specific criteria in each plot. In the original plot, a padj cutoff of 0.05 was used as a threshold for blue and red coloring; log_2_FC less than or equal to −1 were indicated in blue and *log*_2_FC greater than or equal to +1 were indicated in red. Volcano plots that highlighted specific genes used the same cutoffs for light blue and pink, respectively. Highlighted genes were given the same coloring as the original: blue or red.

### Predicted EC Number Assignment

Enzyme commission (EC) numbers can be used to describe both a chemical reaction, and a gene or enzyme that catalyzes that reaction; they are assigned manually using experimental evidence (*116*). EC numbers cannot be officially assigned to the genes in this study, but they can be predicted by comparing genes to well-characterized references, like those in Swiss-Prot: a manually-curated subset of the UniProtKB database (*117*).

Transcripts from the transcriptome assembly were queried against the 2024_06 of 27-Nov-2024 Swiss-Prot release (*117, 118*) using makeblastdb and blastx from the NCBI BLAST+ suite v2.16.0+ (*106*). These queries were performed with default BLAST v2.16.0+ behaviors, but limited the maximum number of high-scoring segment pairs (HSPs) to 1 (-max_hsps 1), and used the following output format: -outfmt “6 qseqid sseqid qlen slen qstart qend sstart send length nident evalue”.

These tabular BLAST results were passed to a series of custom Python and bash scripts that identified query-subject (transcript-protein reference) pairs with the following qualities: amino acid query cover (qcover_aa) of greater than or equal to 40%, subject cover (scover) of greater than or equal to 60%, and alignment percent identity (pident) of greater than or equal to 50%. Several values were used in the calculation of these metrics: the number of identical amino acids in the alignment (nident); the maximum possible amino acid length of the query (max_qlen_aa), defined here as the nucleotide length of the transcript, divided by 3, and rounded down; the number of amino acids included in the alignment itself for both the query and the subject (qalign_len and salign_len, respectively); and the length of the subject (slen).

Using the above variables, qcover_aa and scover were calculated as:

qcover_aa = qalign_len / max_qlen_aa * 100
scover = salign_len / slen * 100

EC numbers, which are annotated in some Swiss-Prot entries, were retrieved using custom Python scripts via UniProt’s REST API (*117, 119*). Transcript queries in query-subject pairs that passed the qcover_aa, scover, and pident filtering steps were given predicted EC numbers, as annotated in their Swiss-Prot subjects. Genes therefore inherited one to several unique predicted EC numbers from their transcripts; genes with multiple predicted EC numbers may catalyze multiple types of reactions. These predicted EC numbers were used to guide downstream analyses, including alignment generation, feature annotation, and metabolic pathway visualization. The functions of some proteins used in volcano plots (like carboxylic ester hydrolases) were identified manually via comparisons made to existing proteins. These manually-identified proteins were not assigned EC numbers primarily because they did not pass the strict 50% identity threshold necessary for assignment. They were thus not included in any downstream analysis that relied on EC number assignment.

### KEGG Pathway Visualization

The Kyoto Encyclopedia of Genes and Genomes (KEGG) includes a database of manually-drawn metabolic pathways (*71–73*). These pathways include both EC numbers and small molecule identifiers; genes that were assigned EC numbers and small molecules that were identified during mass spectrometry steps could therefore be mapped to these pathways. Expression data was projected onto the KEGG pathway for flavonoid biosynthesis (map00941) in two different ways, depending on data type.

KEGG EC-gene expression: Following the assignment of predicted EC numbers to individual genes, each unique EC number could be represented by at least one gene and its expression information. Instead of selecting a single gene to represent each EC number, *log*_2_FC were averaged across all genes that shared a particular EC number. The average of *log*_2_FC is equivalent to the *log*_2_ geometric mean of the FC. Only genes whose padj were less than 0.05 were included in this calculation, thus limiting *log*_2_FC representation to genes that were hypothesized to be significantly differentially expressed in JA-treated samples when compared with water-treated controls. These *log*_2_FC averages were initially projected onto the KEGG flavonoid pathway using KEGG’s online Color tool, with an inclusive lower limit of −1 (blue), and an inclusive upper limit of +1 (red). This represents a treated/control FC of 1/2x and 2x, respectively. Average *log*_2_FC that exceeded these limits were visually represented as the limits themselves. EC numbers that were not assigned to significantly differentially expressed genes were colored grey; EC numbers that were not assigned to any gene in the transcriptome assembly were left white.

KEGG small molecule expression: Small molecules, represented by circles, and labeled by numbers, were colored according to two different FC comparisons: JA-fed / unfed control, and JA-unfed / unfed control. An inclusive lower limit of −0.5 *log*_2_FC (cyan), and an inclusive upper limit of +0.5 *log*_2_FC (magenta) were used to visualize expression in both groups. Initial projections of these data were generated using KEGG’s online Color tool, and combined into a final figure. JA-unfed *log*_2_FCs were represented by solid fill colors inside small molecule circles, while JA-fed *log*_2_FCs were represented by larger, transparent circles, centered on original small molecule circles. The flavonoid pathway itself was manually recreated in Inkscape (https://inkscape.org/), and all data were incorporated in the final figure. A table of small molecule names, according to number, can be found in the Supplementary Information (SI).

### Alignments and Feature Annotation

Protein coding sequences were identified manually after translating transcripts; the start sites of some proteins were extended to the 5’ ends of their transcripts if either no explicit start codon could be identified, or if there was no explicit stop codon between the 5’ end and the first start codon. Features like active site residues, which were annotated and used as a basis for inference throughout this work, were largely identified using annotated alignments generated with Computer-Assisted Sequence Annotation (CASA), downloaded from https://github.com/grtakaha/CASA in May, 2025 (*38*).

### Genome Annotation

GCA_001925005.1_ASM192500v1_genomic.fna was unmasked using an awk script from NCBI’s site (https://www.ncbi.nlm.nih.gov/genome/doc/ftpfaq/#masking). It was then soft-masked with RepeatMasker version open-4.0.7 (*120*) using only repeats from the *Drosera capensis* entry in the Plant Repeat Database (PlantRep, http://www.plantrep.cn/, (*121*)). This soft-masked genome was annotated using the MAKER v3.01.04 pipeline (*122–124*) in a series of steps adapted from (*33, 122, 125*) and https://darencard.net/blog/2017-05-16-maker-genome-annotation/. Notably, trna and keep_preds were set to 0 in control files for all steps.

MAKER was first run using the TSA-submitted transcriptome (GKUK00000000) and a curated set of UniProt (*117*) proteins as evidence; the UniProt set included all Caryophyllales proteins that were accessible on 2024.03.01, with evidence at the transcript or protein level. Gene models whose annotation edit distances (AED) were 0.25 or less, and whose protein coding regions contained at least 50 amino acids were saved separately and used to train SNAP (downloaded from https://github.com/KorfLab/SNAP on April 25, 2024) (*126*), one of several ab initio gene predictors employed in the MAKER pipeline. Training was followed by another round of MAKER annotations, this time with SNAP predictions included. Confident gene models from this round of annotations were then used to train SNAP once more. MAKER was run one final time using the D. capensis transcriptome, the UniProt Caryophyllales proteins, and the most recently-trained SNAP models, along with two alternate sources of evidence (the Dionaea muscipula transcriptome (*127*), the Nepenthes mirabilis transcriptome (*128*)), and one other ab initio predictor Augustus v3.2.3 (*129*) (using the built-in tomato HMM).

In summary, the MAKER steps used in this study followed the following schema: annotation (transcript, protein), SNAP training, annotation (transcript, protein, SNAP), SNAP training, final annotation (transcript, protein, SNAP, two alternate transcripts, Augustus – tomato).

Only gene models from MAKER itself (rather than individual predictors or evidence) were retained for downstream applications in the final annotation. These were gene models with “maker” in column 2 of the GFF3-formatted annotation file.

This final MAKER annotation had 27813 genes with an average length of 3862.54 bp. 95.6% of these genes had an AED of 0.5 or less, based on the given transcriptome and protein evidence. The BUSCO v5.7.1 score for mRNA transcripts from this set of annotations was 82.6% (complete BUSCOs), using the embryophyta dataset (embryophyte_odb10), which contains 1614 BUSCOs.

Functional annotations were added to GFF3, transcript FASTA, and protein FASTA files using a combination of InterProScan v5.64-96.0 (*130*) and BLAST v2.15.0 (*106*), according to support protocol 3 and basic protocol 5 from Campbell et al. (*122*). Swiss-Prot release 2024_02 of 27-Mar-2024 (*117, 118*) was used as the subject database for BLAST. Additional qualifiers (protein_id, transcript_id, locus_tag, and product) were added to column 9 of the GFF3 file in preparation for submission to the NCBI’s WGS database. Product qualifiers were added based on MAKER’s functional annotations and NCBI guidelines. A .sqn file was generated for submission using table2asn (https://www.ncbi.nlm.nih.gov/genbank/table2asn/) using the final GFF3 file and an edited version of the original NCBI genome (GCA_001925005.1_ASM192500v1_genomic.fna). On recommendation from NCBI, scaffolds in this version of the genome were renamed, and several newly-identified duplicate or contaminant scaffolds were removed. This submission can be found under the LIEC00000000 ID.

---

**Table S1:** Previously published aspartic proteases from *D. capensis* found in the transcriptome. Proteins with no published ID were in the original genome (*33*) annotation, but were not used in published analyses. Only *log*_2_FC greater than or equal to 1 or less than or equal to −1 were assigned “up” and “down” changes, respectively.

| transcriptome | genome | published ID | name | change | reference |
| --- | --- | --- | --- | --- | --- |
| DCAP.26675.1 | AKJ16_0003055-RA |  |  | up | (33) |
| DCAP.9785.1 | AKJ16_0007732-RA |  |  | up | (33) |
| DCAP.9784.1 | AKJ16_0007106-RA | DCAP_7106 | NEP_DCAP | up | (33) |
| DCAP.12239.1 | AKJ16_0007106-RA | DCAP_7106 | NEP_DCAP | none | (33) |
| DCAP.18786.1 | AKJ16_0004149-RA | DCAP_4149 | Droserasin 1 | none | (33) |
| DCAP.27390.1 | AKJ16_0003196-RA | DCAP_3196 | Droserasin 2 | none | (33) |
| DCAP.23195.2 | AKJ16_0004536-RA |  |  | none | (33) |

**Table S2:**
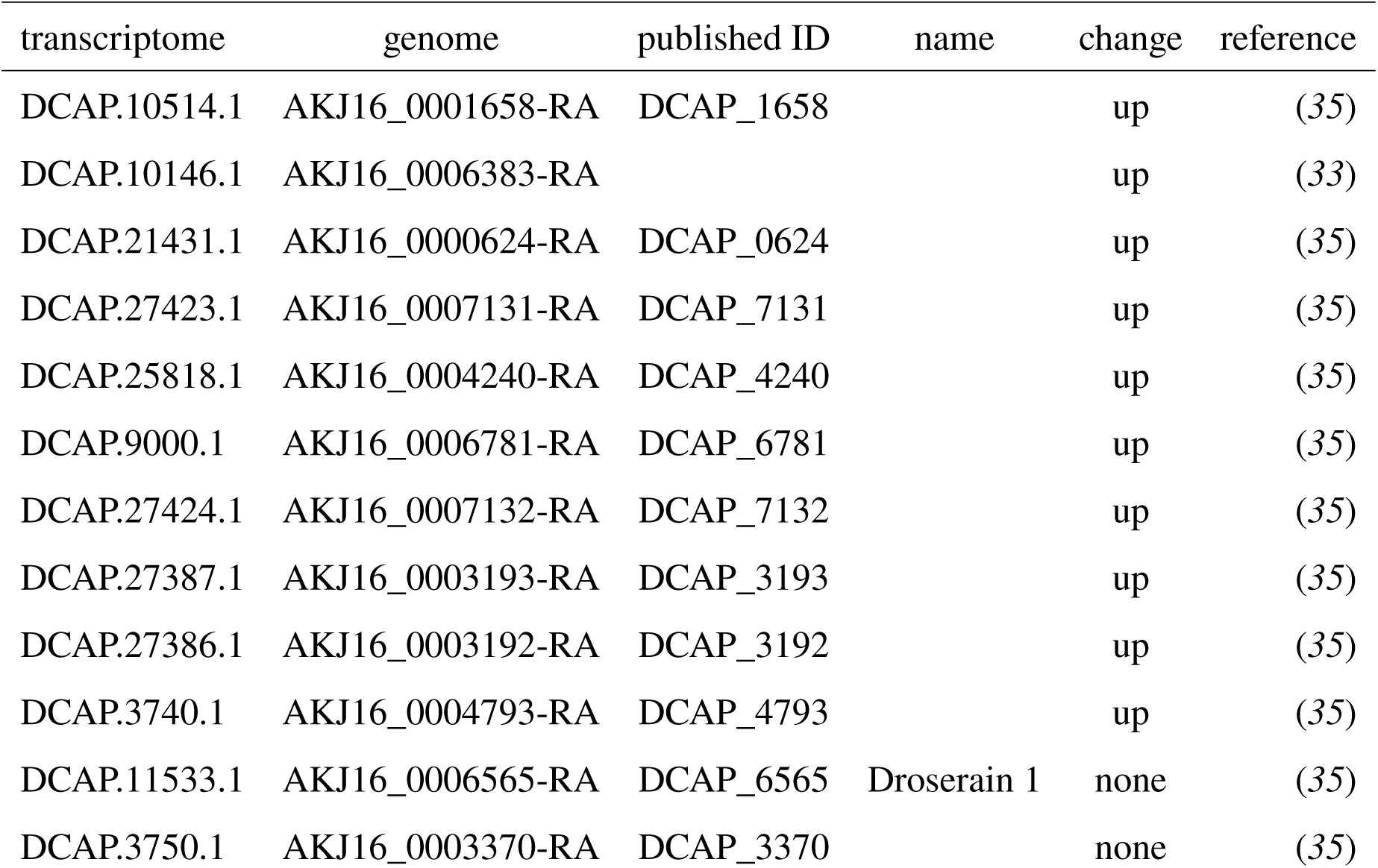
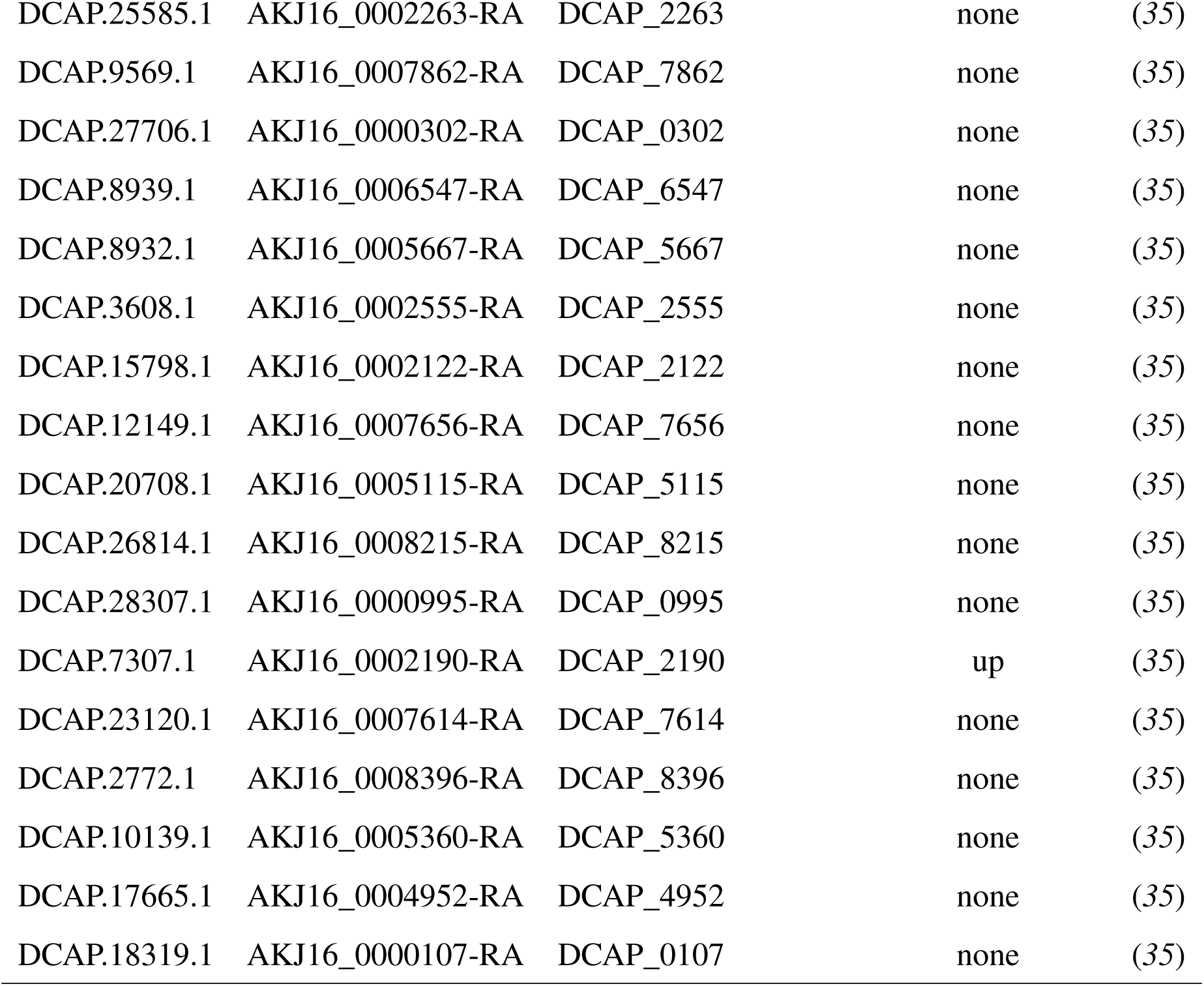
Previously published cysteine proteases from *D. capensis* found in the transcriptome. Proteins with no published ID were in the original genome (*33*) annotation, but were not used in published analyses. Only *log*_2_FC greater than or equal to 1 or less than or equal to −1 were assigned “up” and “down” changes, respectively.

| transcriptome | genome | published ID | name | change | reference |
| --- | --- | --- | --- | --- | --- |
| DCAP.10514.1 | AKJ16_0001658-RA | DCAP_1658 |  | up | (35) |
| DCAP.10146.1 | AKJ16_0006383-RA |  |  | up | (33) |
| DCAP.21431.1 | AKJ16_0000624-RA | DCAP_0624 |  | up | (35) |
| DCAP.27423.1 | AKJ16_0007131-RA | DCAP_7131 |  | up | (35) |
| DCAP.25818.1 | AKJ16_0004240-RA | DCAP_4240 |  | up | (35) |
| DCAP.9000.1 | AKJ16_0006781-RA | DCAP_6781 |  | up | (35) |
| DCAP.27424.1 | AKJ16_0007132-RA | DCAP_7132 |  | up | (35) |
| DCAP.27387.1 | AKJ16_0003193-RA | DCAP_3193 |  | up | (35) |
| DCAP.27386.1 | AKJ16_0003192-RA | DCAP_3192 |  | up | (35) |
| DCAP.3740.1 | AKJ16_0004793-RA | DCAP_4793 |  | up | (35) |
| DCAP.11533.1 | AKJ16_0006565-RA | DCAP_6565 | Droserain 1 | none | (35) |
| DCAP.3750.1 | AKJ16_0003370-RA | DCAP_3370 |  | none | (35) |
| DCAP.25585.1 | AKJ16_0002263-RA | DCAP_2263 |  | none | (35) |
| DCAP.9569.1 | AKJ16_0007862-RA | DCAP_7862 |  | none | (35) |
| DCAP.27706.1 | AKJ16_0000302-RA | DCAP_0302 |  | none | (35) |
| DCAP.8939.1 | AKJ16_0006547-RA | DCAP_6547 |  | none | (35) |
| DCAP.8932.1 | AKJ16_0005667-RA | DCAP_5667 |  | none | (35) |
| DCAP.3608.1 | AKJ16_0002555-RA | DCAP_2555 |  | none | (35) |
| DCAP.15798.1 | AKJ16_0002122-RA | DCAP_2122 |  | none | (35) |
| DCAP.12149.1 | AKJ16_0007656-RA | DCAP_7656 |  | none | (35) |
| DCAP.20708.1 | AKJ16_0005115-RA | DCAP_5115 |  | none | (35) |
| DCAP.26814.1 | AKJ16_0008215-RA | DCAP_8215 |  | none | (35) |
| DCAP.28307.1 | AKJ16_0000995-RA | DCAP_0995 |  | none | (35) |
| DCAP.7307.1 | AKJ16_0002190-RA | DCAP_2190 |  | up | (35) |
| DCAP.23120.1 | AKJ16_0007614-RA | DCAP_7614 |  | none | (35) |
| DCAP.2772.1 | AKJ16_0008396-RA | DCAP_8396 |  | none | (35) |
| DCAP.10139.1 | AKJ16_0005360-RA | DCAP_5360 |  | none | (35) |
| DCAP.17665.1 | AKJ16_0004952-RA | DCAP_4952 |  | none | (35) |
| DCAP.18319.1 | AKJ16_0000107-RA | DCAP_0107 |  | none | (35) |

**Table S3:** Previously published chitinases from *D. capensis* found in the transcriptome. Proteins with no published ID were in the original genome (*33*) annotation, but were not used in published analyses. Only *log*_2_FC greater than or equal to 1 or less than or equal to −1 were assigned “up” and “down” changes, respectively.

| transcriptome | genome | published ID | type | change | reference |
| --- | --- | --- | --- | --- | --- |
| DCAP.77.1 | AKJ16_0007544-RA | DCAP_7544 | Family 18 | up | (36) |
| DCAP.13060.1 | AKJ16_0002209-RA | DCAP_2209 | Family 18 | up | (36) |
| DCAP.18317.1 | AKJ16_0000106-RA | DCAP_0106 | Family 18 | up | (36) |
| DCAP.21069.1 | AKJ16_0004817-RA | DCAP_4817 | Family 19 | up | (36) |
| DCAP.21889.1 | AKJ16_0004817-RA | DCAP_4817 | Family 19 | up | (36) |
| DCAP.21847.1 | AKJ16_0005455-RA | DCAP_5455 | Family 18 | none | (36) |
| DCAP.12510.1 | AKJ16_0002737-RA | DCAP_2737 | Family 18 | none | (36) |

**Table S4:** Previously published esterase/lipases from *D. capensis* found in the transcriptome. Proteins with no published ID were in the original genome (*33*) annotation, but were not used in published analyses. Only *log*_2_FC greater than or equal to 1 or less than or equal to −1 were assigned “up” and “down” changes, respectively.

| transcriptome | genome | published ID | change | reference |
| --- | --- | --- | --- | --- |
| DCAP.23163.1 | AKJ16_0005138-RA | DCAP_5138 | up | (37) |
| DCAP.16632.1 | AKJ16_0001301-RA |  | up | (33) |
| DCAP.7535.1 | AKJ16_0008291-RA |  | up | (33) |
| DCAP.22864.1 | AKJ16_0004076-RA | DCAP_4076 | down | (37) |
| DCAP.6000.1 | AKJ16_0005759-RA |  | down | (33) |
| DCAP.11566.1 | AKJ16_0002838-RA |  | down | (33) |
| DCAP.11134.1 | AKJ16_0006666-RA |  | down | (33) |
| DCAP.22778.1 | AKJ16_0005587-RA | DCAP_5587 | down | (37) |
| DCAP.4991.1 | AKJ16_0006947-RA | DCAP_6947 | none | (37) |
| DCAP.22003.1 | AKJ16_0003343-RA | DCAP_3343 | none | (37) |
| DCAP.668.1 | AKJ16_0004098-RA | DCAP_4098 | none | (37) |
| DCAP.7299.1 | AKJ16_0002187-RA | DCAP_2187 | none | (37) |
| DCAP.8086.1 | AKJ16_0006217-RA | DCAP_6217 | none | (37) |
| DCAP.7571.1 | AKJ16_0001761-RA | DCAP_1761 | none | (37) |

**Table S5:** Small molecules in the flavonoid biosynthesis pathway.

| number | name |
| --- | --- |
|  | Cinnamoyl-CoA |
| 1. |  |
| 2. | Pinocembrin chalcone |
| 3. | Pinocembrin |
| 4. | Pinobanksin |
| 5. | Pinobanksin 3-acetate |
| 6. | Chrysin |
| 7. | Pinostrobin |
| 8. | Galangin |
| 9. | Isoliquiritigenin |
| 10. | Liquiritigenin |
| 11. | Garbanzol |
| 12. | 5-Deoxyleucopelargonidin |
| 13. | 7,4'-Dihydroxyflavone |
| 14. | Butein |
| 15. | Butin |
| 16. | Dihydrofisetin |
| 17. | Deoxyleucocyanidin |
| 18. | Dihydro-4-coumaroyl-CoA |
| 19. | Phloretin |
| 20. | Phlorizin |
| 21. | Prunin |
| 22. | Naringin |
| 23. | Desmethyl-xanthohumol |
| 24. | Xanthohumol |
| 25. | 4-O-Methyl-xanthohumol |
| 26. | Hesperetin |

**Table S5: Small molecules in the flavonoid biosynthesis pathway.**
| number | name |
| --- | --- |
| 27. | Hesperetin 7-O-glucoside |
| 28. | Neohesperidin |
| 29. | Isosakuranetin |
| 30. | p-Coumaroyl-CoA |
| 31. | Naringenin chalcone |
| 32. | Naringenin |
| 33. | Dihydrokaempferol |
| 34. | Leucopelargonidin |
| 35. | Pelargonidin |
| 36. | (-)-Epiafzelechin |
| 37. | Sakuranetin |
| 38. | p-Coumaroyl shikimic acid |
| 39. | p-Coumaroyl quinic acid |
| 40. | Naringenin chalcone 4'-O-glucoside |
| 41. | Aureusidin 6-O-glucoside |
| 42. | Apiforol |
| 43. | (+)-Afzelechin |
| 44. | 8-C-Glucosylnaringenin |
| 45. | Vitexin |
| 46. | Kaempferol |
| 47. | Chalcone 2'-O-glucoside |
| 48. | Apigenin |
| 49. | Caffeoyl shikimic acid |
| 50. | Caffeoyl quinic acid |
| 51. | 2',3,4,4',6'-Peptahydroxychalcone 4'-O-glucoside |
| 52. | Bracteatin 6-O-glucoside |
| 53. | Quercetin |

**Table S5: Small molecules in the flavonoid biosynthesis pathway.**
| number | name |
| --- | --- |
| 54. | Caffeoyl-CoA |
| 55. | 2',3,4,4',6'-Pentahydroxychalcone |
| 56. | Eriodictyol |
| 57. | Dihydroquercetin |
| 58. | Leucocyanidin |
| 59. | Cyanidin |
| 60. | (-)-Epicatechin |
| 61. | Luteolin |
| 62. | (+)-Catechin |
| 63. | Luteoforol |
| 64. | Feruloyl-CoA |
| 65. | 4,2',4',6'-Tetrahydroxy-3-methoxychalcone |
| 66. | Homoeriodictyol |
| 67. | Tricetin |
| 68. | Myricetin |
| 69. | Dihydrotricetin |
| 70. | Dihydromyricetin |
| 71. | Leucodelphinidin |
| 72. | Delphinidin |
| 73. | (-)-Epigallocatechin |
| 74. | (+)-Gallocatechin |

**Figure S1:**
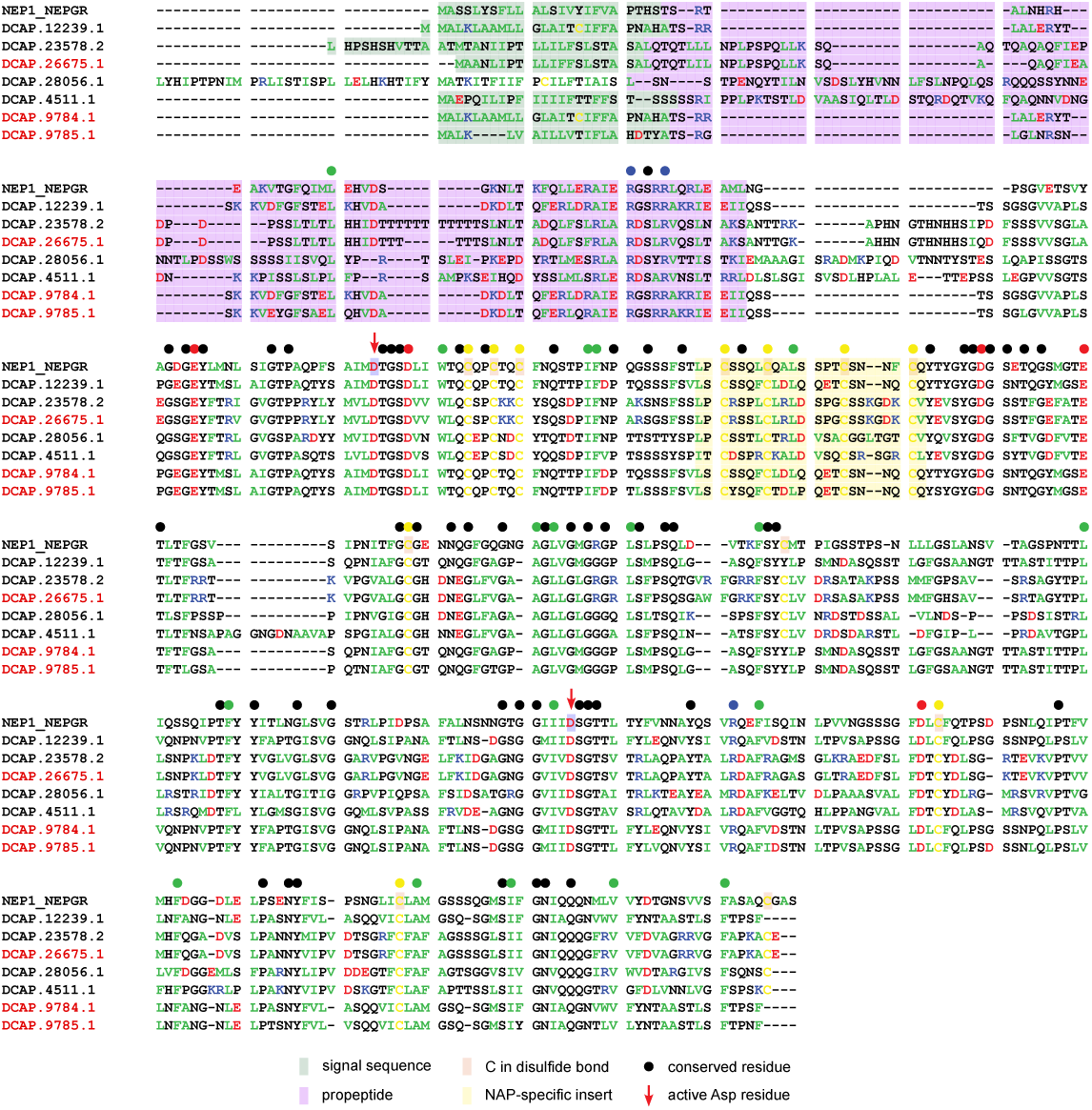
Nepenthesin sequence alignment. These proteins are homologs of Nepenthesin 1 (UniProt ID: NEP1_NEPGR) from *Nepenthes gracilis* (*46*), which is also a carnivorous plant in Caryophylalles. Sequence identifiers refer to the *D. capensis* transcriptome. Labels in red indicate that the protein is upregulated in response to jasmonic acid treatment; those in black indicate no significant change. Closed circles indicate conserved residues, with amino acid properties indicated as follows: positively charged, blue; negatively charged, red; cysteine, yellow; hydrophobic, green; other, black. Active site Asp residues are labeled with red arrows. Signal sequences are highlighted in green. Pro-sequences are highlighted in purple. Cysteines in disulfide bonds (in the reference sequence) are highlighted in light orange. Nepenthesins have a unique sequence feature, the nepenthesin aspartic protease (NAP)-specific insert, which is highlighted in yellow.

**Figure S2:**
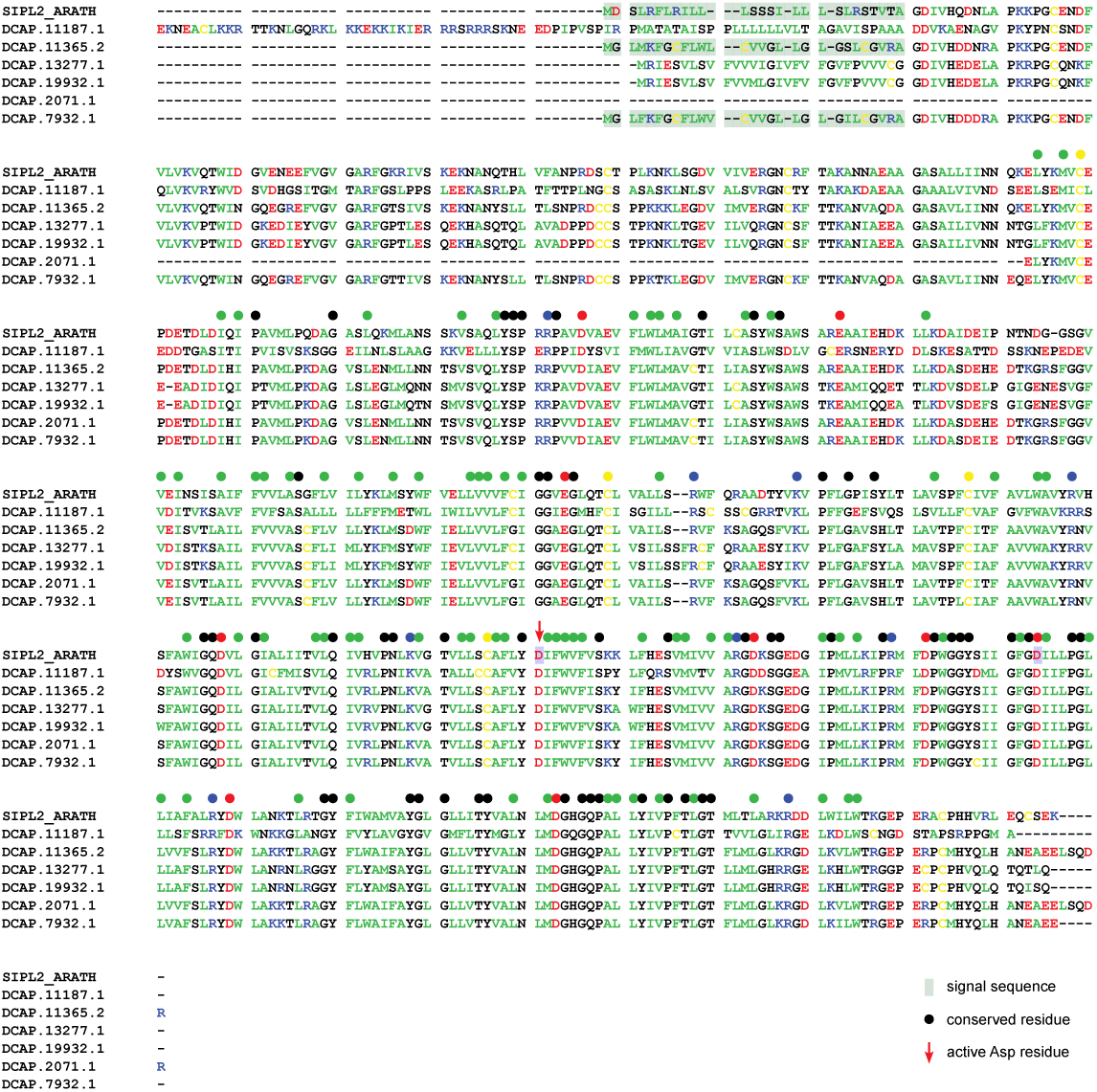
Signal peptidases. SIPL2_ARATH, a signal peptidase-like type 2, is a membrane protein that cleaves signal peptides of other proteins in the membrane (*55, 131*). These proteases are neither up- nor downregulated in response to jasmonic acid. Annotations are as in **Supplementary Figure S1**.

**Figure S3:**
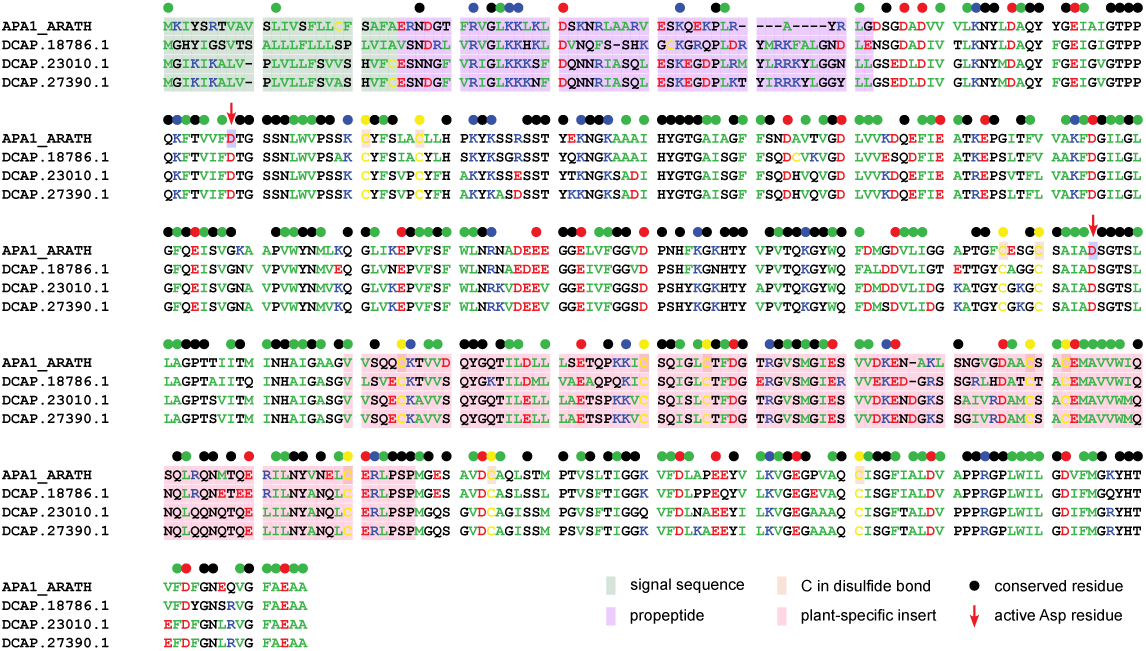
Homologs of APA1_ARATH. APA1_ARATH in *Arabidopsis thaliana* breaks down storage proteins (*51*). *D. capensis* has three paralogs, none of which is upregulated in response to jasmonic acid. These proteases have a unique 100-residue feature called the plant-specific insert (PSI), which is cleaved from the mature protein and acts as an independent anti-microbial peptide (*132*). Here, the PSI is highlighted in pink. Annotations are otherwise as in **Supplementary Figure S1**.

**Figure S4:**
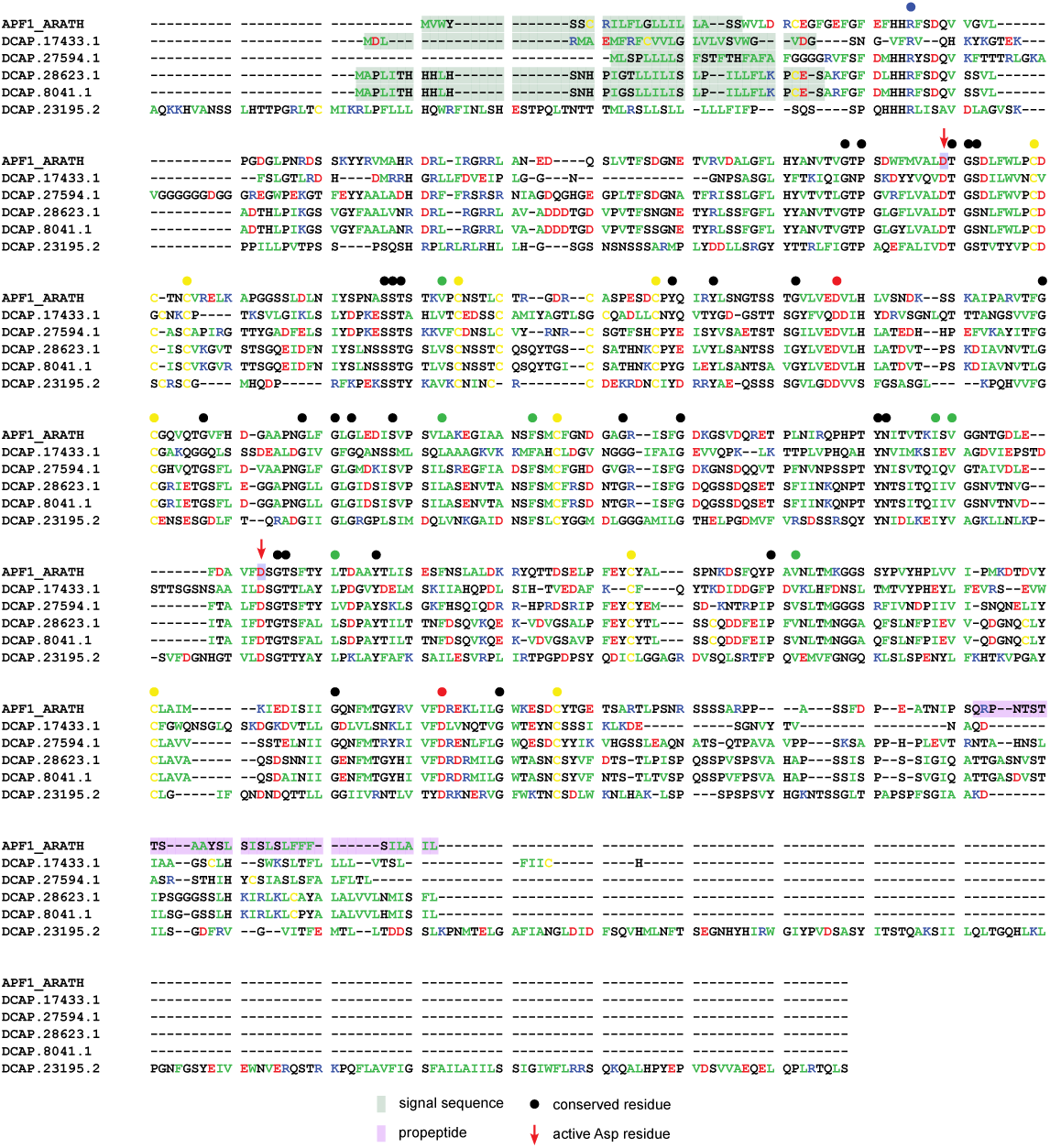
Homologs of APF1_ARATH. *D. capensis* has five paralogs of APF1_ARATH (*131*), none of which is upregulated in response to jasmonic acid. Annotations are as in **Supplementary Figure S1**.

**Figure S5:**
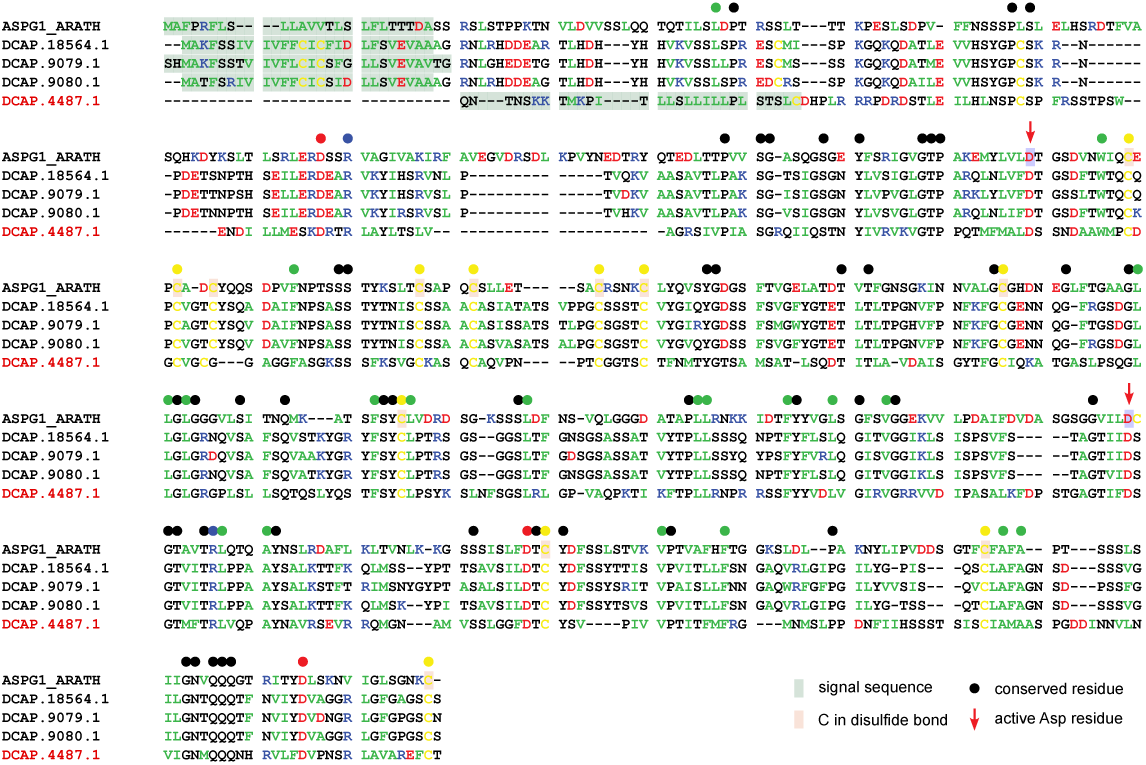
Homolog of ASPG1_ARATH. ASPG1_ARATH is Aspartic Protease in Guard Cell 1. It is involved in drought response and is regulated by the abscisic acid pathway (*49*). *D. capensis* has four homologs, one of which is upregulated in response to jasmonic acid. Annotations are as in **Supplementary Figure S1**.

**Figure S6:**
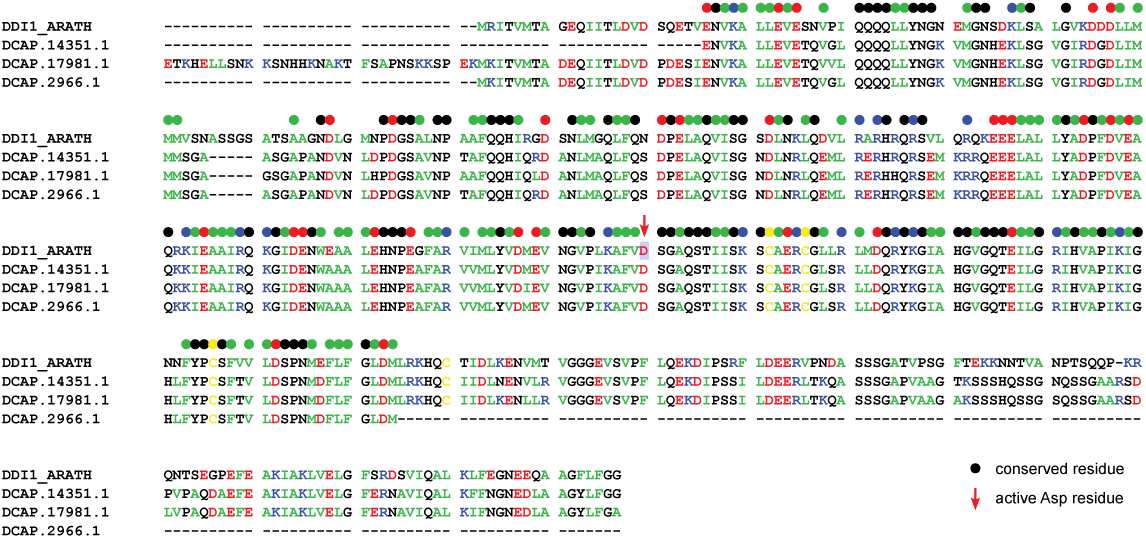
DDI1_ARATH. DNA-Damage Inducible 1 is involved in degradation of ubiquitinated proteins in *Arabidopsis thaliana* (*54*). Annotations are as in **Supplementary Figure S1**.

**Figure S7:**
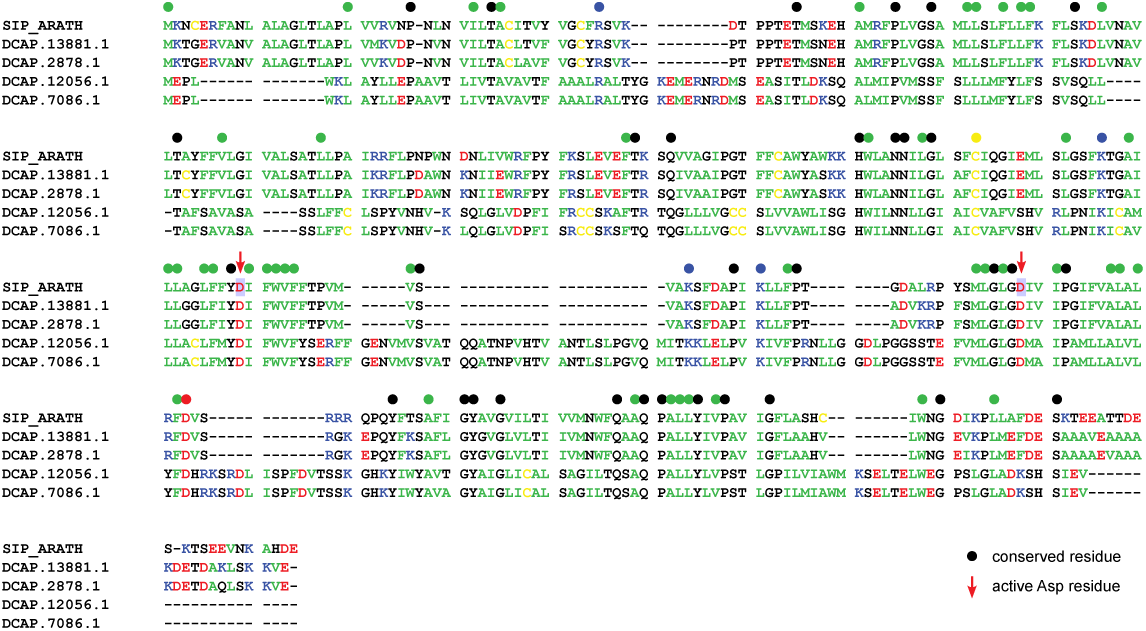
SIP_ARATH. This signal peptide peptidase is required for pollen development (*56*). It is a membrane protein with nine transmembrane helices. Annotations are as in **Supplementary Figure S1**.

**Figure S8:**
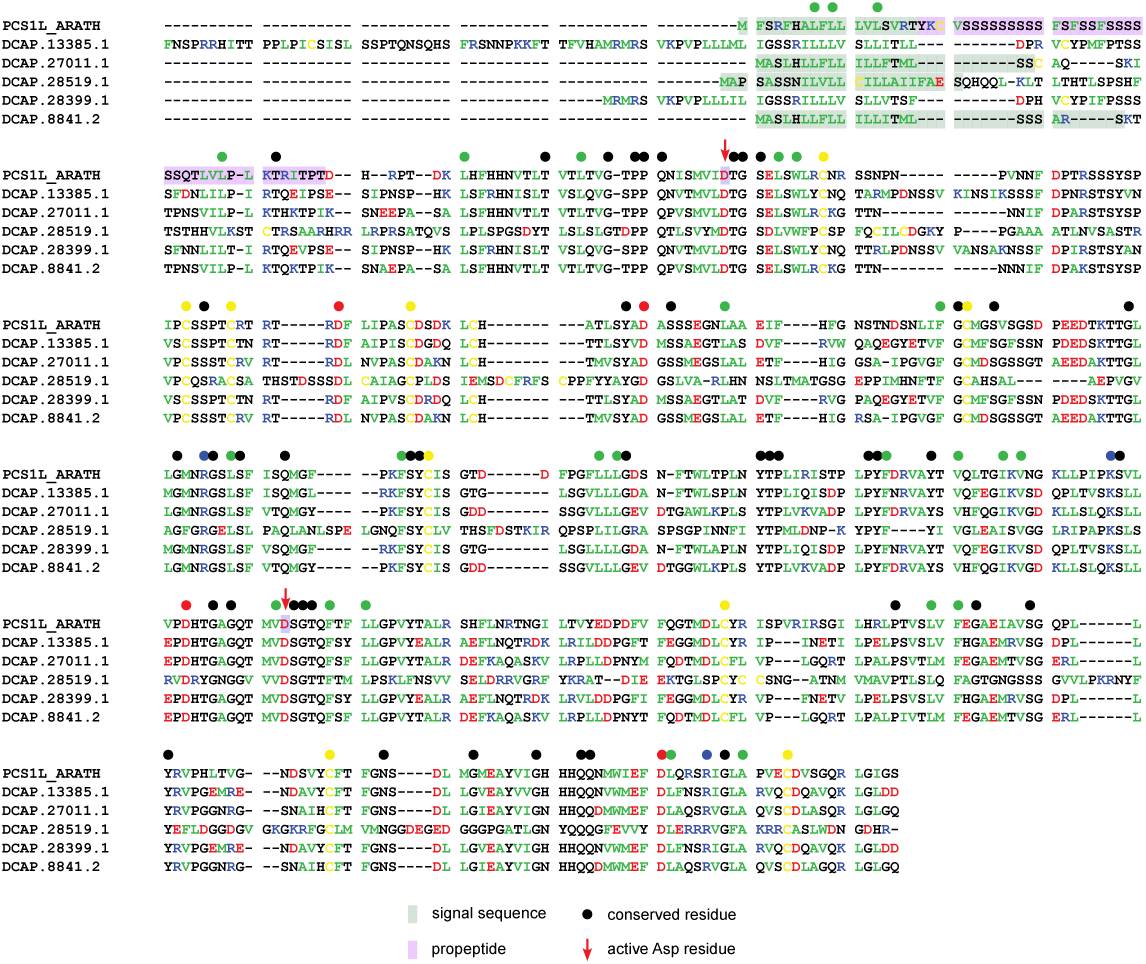
PCS1L_ARATH. In *Arabidopsis*, this protein suppresses programmed cell death during embryonic development (*57*). Annotations are as in **Supplementary Figure S1**.

**Figure S9:**
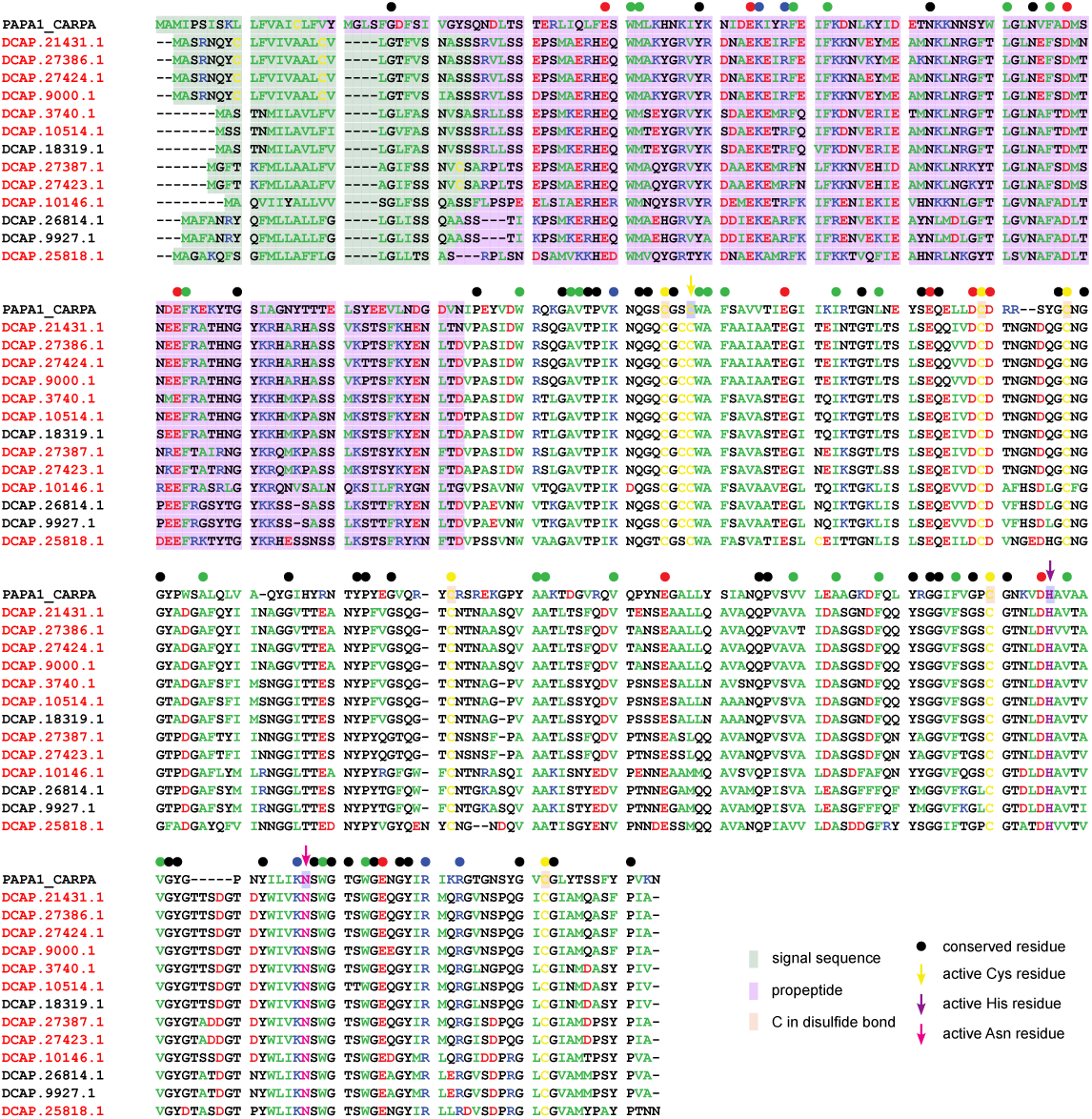
Cysteine proteases. *D. capensis* has many cysteine proteases. Thirteen are shown here aligned with papain from *Carica papaya* (UniProt ID: PARA1_CARPA) The active site is characterized by a catalytic triad consisting of Cys (yellow arrow), His (purple arrow), and Asn (magenta arrow). Annotations are otherwise as in **Supplementary Figure S1**.

**Figure S10:**
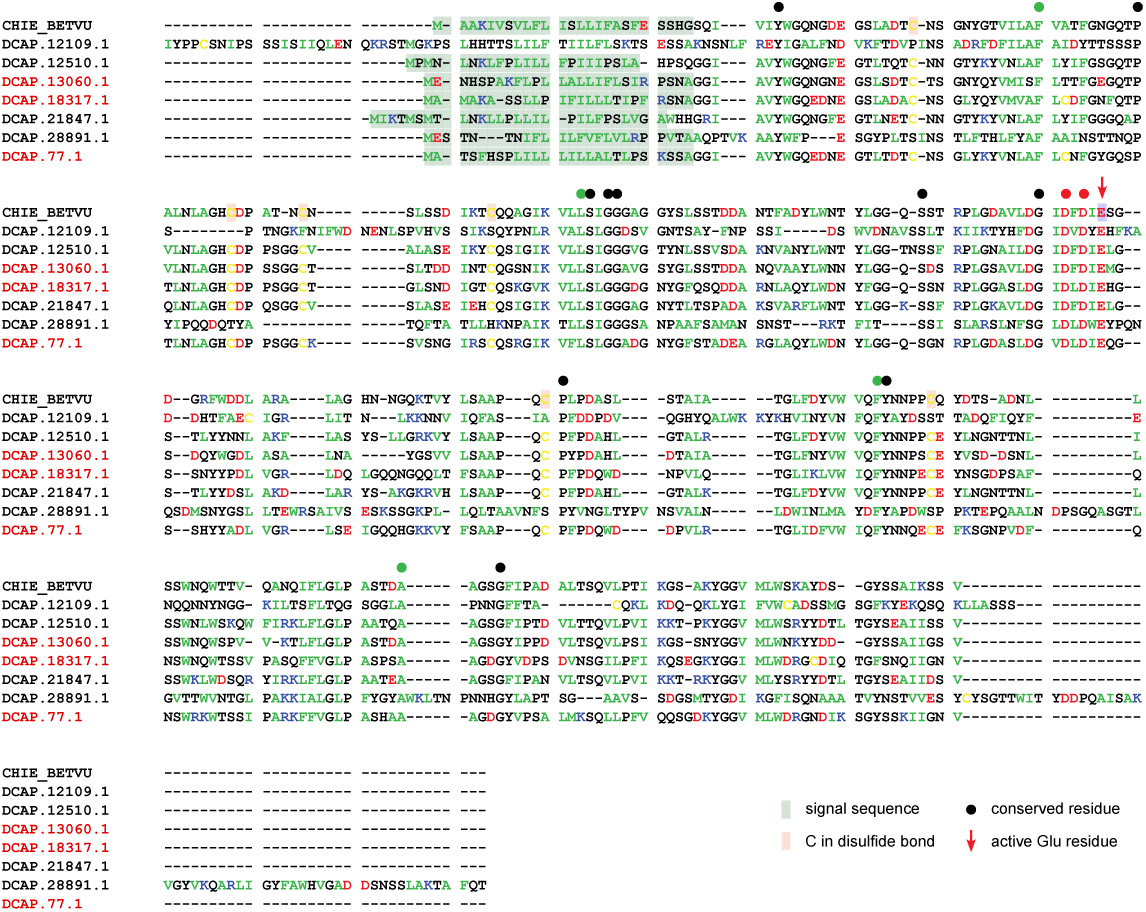
Family 18 Chitinases. Here we identify seven paralogs of Family 18 chitinases from *Drosera capensis*. They are annotated by comparison to Acidic endochitinase SE2 (UniProt ID CHIE_BETVU) (*133*). Three are upreglated in response to jasmonic acid, while four remain unchanged. The catalytic residue is a Glu, indicated with a red arrow. Annotations are otherwise as in **Supplementary Figure S1**.

**Figure S11:**
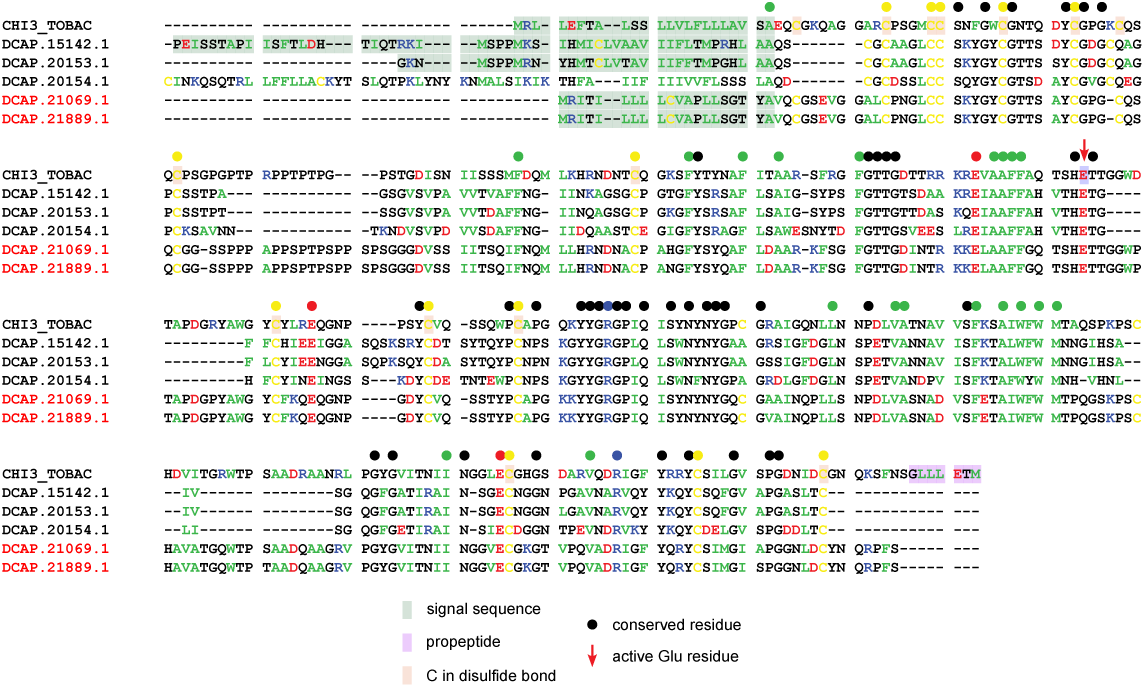
Family 19 chitinases. Five Family 19 chitinase paralogs were found in *D. capensis*. Two are upregulated in response to jasmonic acid, while three are unchanged. Here, annotations based on Endochitinase 3 from *Nicotiana tabacum* (Uniprot ID: CHI3_TABAC), which is involved in defense against fungal pathogens (*66*). Annotations are as in **Supplementary Figure S10**.

**Figure S12:**
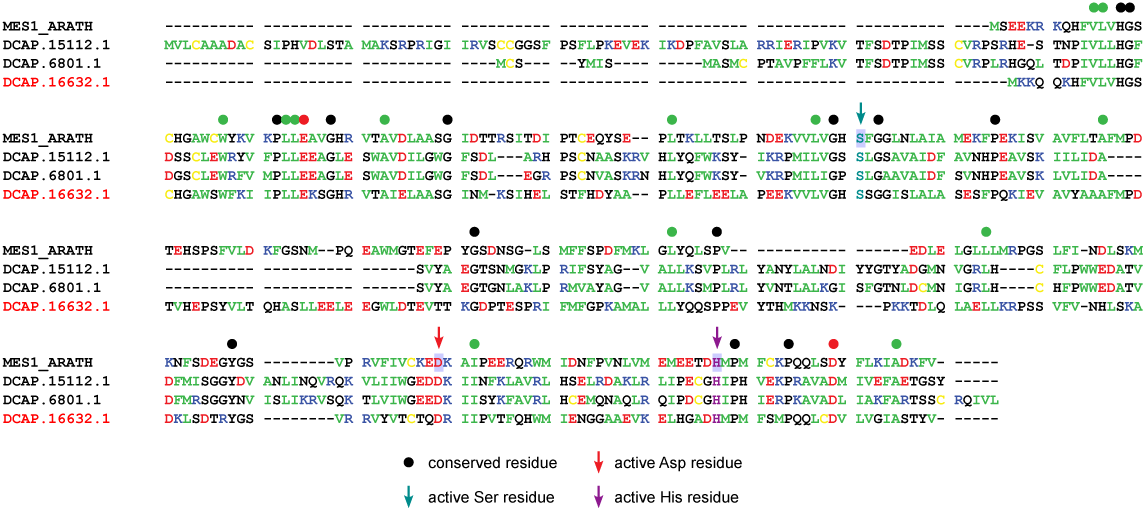
Methylesterases. *D. capensis* has three methylesterases homologous to UniProt ID: MES1_ARATH from *Arabidopsis thaliana*. One of these is upregulated and two are unchanged in response to jasmonic acid treatment. The *Arabidopsis* homolog has carboxylesterase, methyl indole-3-acetic acid esterase, methyl salicylate esterase activity, and methyl jasmonate esterase activity (*134*). These proteins have a catalytic triad consisting of Ser (dark cyan arrow), His (purple arrow), and Asp (red arrow). Annotations are otherwise as in **Supplementary Figure S1**.

**Figure S13:**
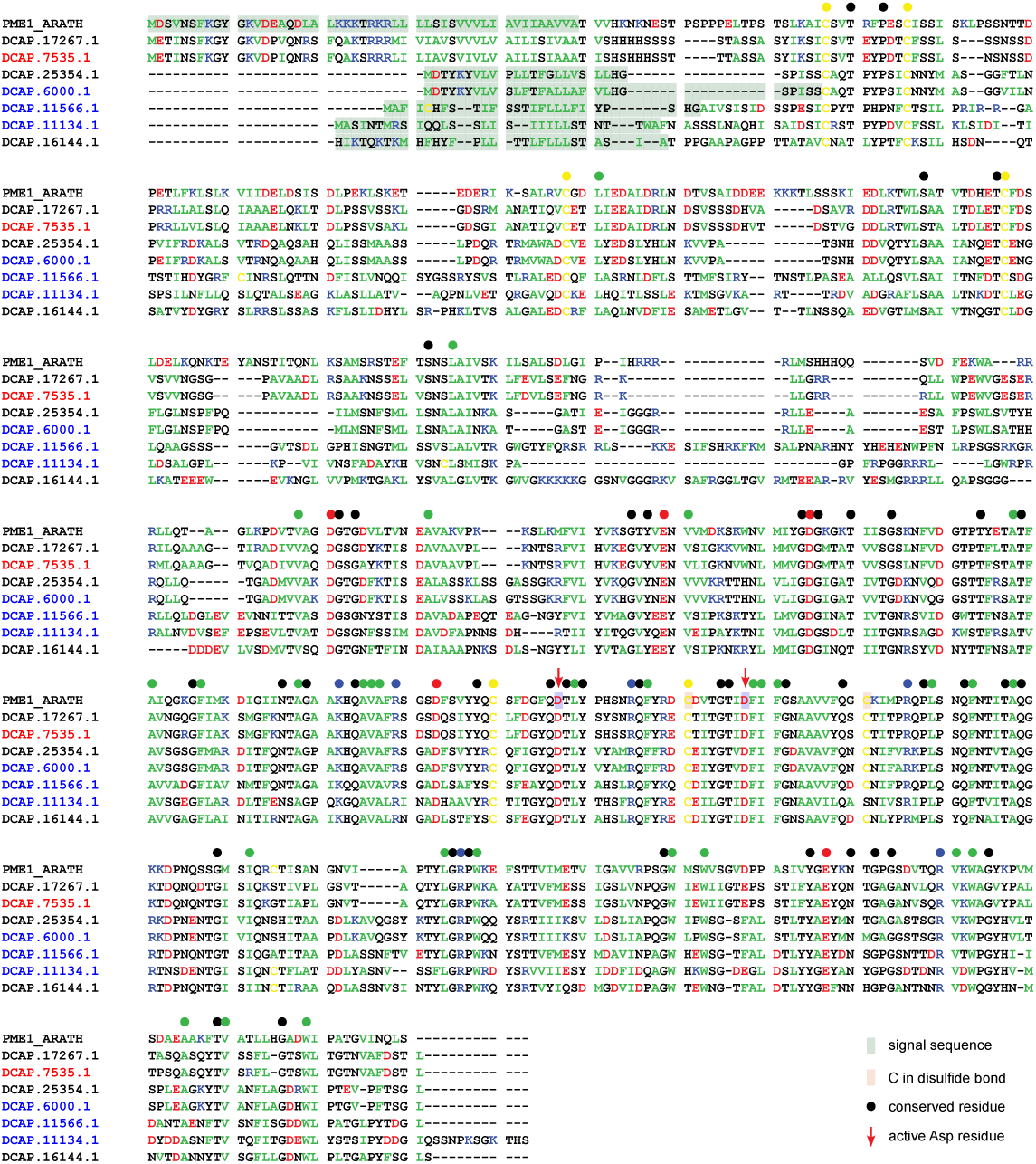
Pectinesterases. *D. capensis* has seven homologs of *Arabidopsis* Pectinesterase 1 (UniProt ID: PME1_ARATH). This protein modifies cell walls by demethylesterification of pectin (*131*). Of these, one is up-regulated and three are downregulated in response to jasmonic acid treatment. These proteins have two catalytic Asp residues (red arrows). Annotations are otherwise as in **Supplementary Figure S1**.

**Figure S14:**
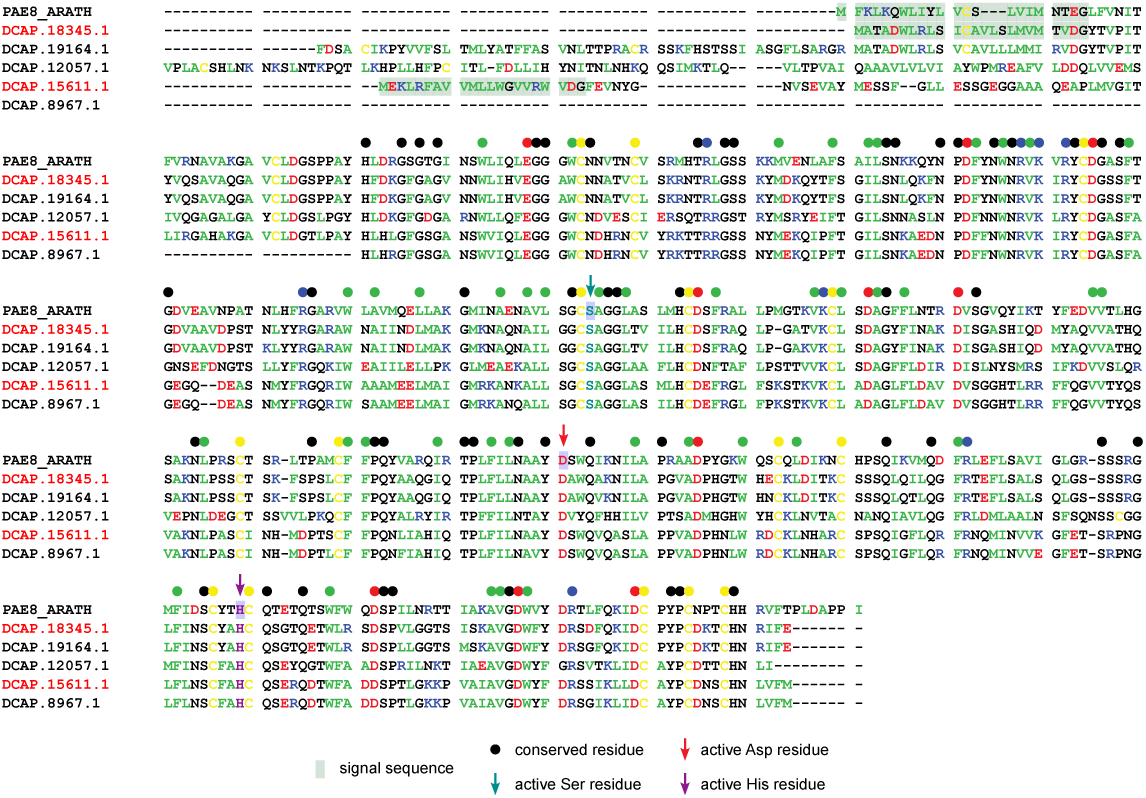
Pectin acetylesterases. *D. capensis* has five homologs of *Arabidopsis thaliana* Pectin acetylesterase 8 (UniProt ID: PAE8_ARATH), an enzyme that modifies cell wall pectin by hydrolyzing acetyl esters in homogalacturonan regions (*135*). Two of these are upregulated in response to jasmonic acid treatment. These proteins have a catalytic triad consisting of Ser (dark cyan arrow), His (purple arrow), and Asp (red arrow). Annotations are otherwise as in **Supplementary Figure S1**.

**Figure S15:**
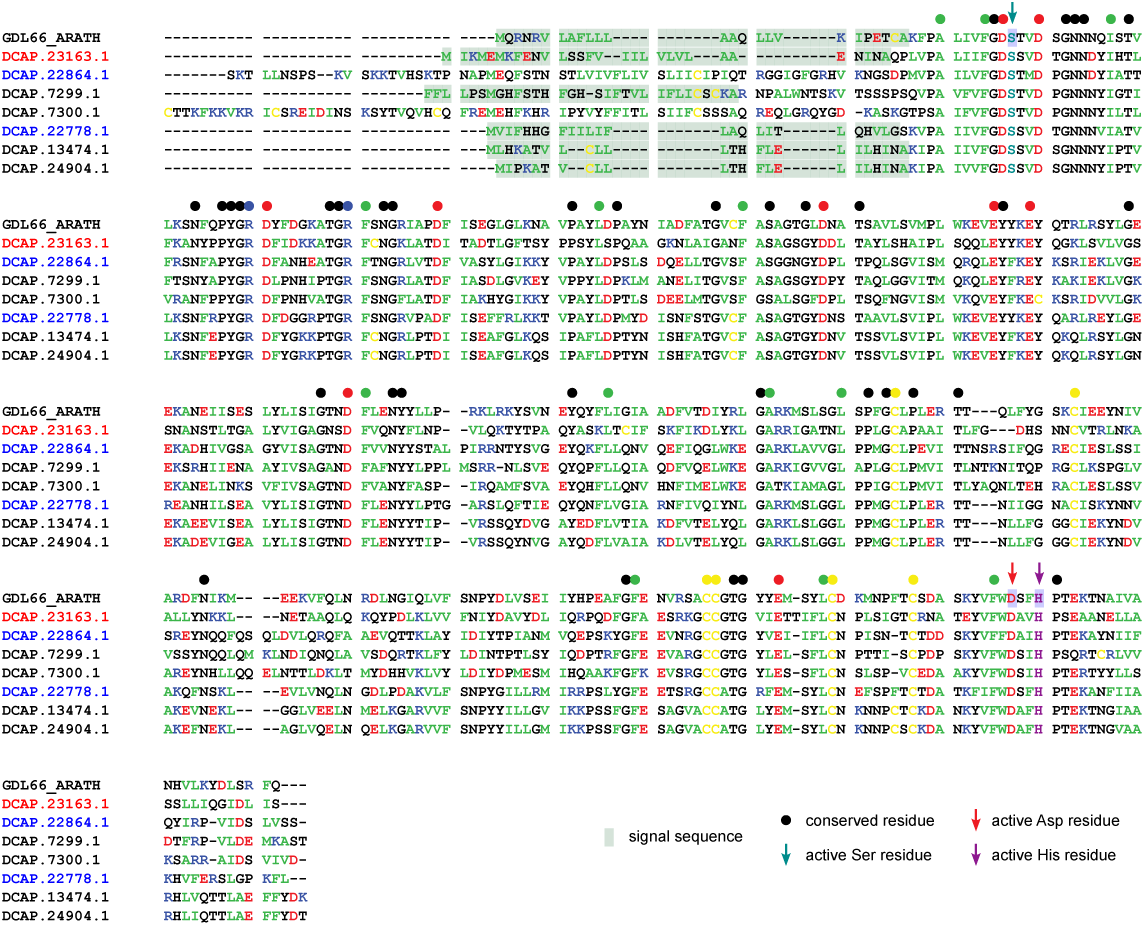
GDSL esterase/lipases. *D. capensis* has seven homologs of *Arabidopsis* GDSL esterase/lipase (UniProt ID: GDL66_ARATH). GDSL esterase/lipases have a broad range of activity and substrate specificity, ranging from hydrolyze esters or acyl esters to forming polyester compounds under some conditions (*136, 137*). Of these, one is upregulated and two are downregulated in response to jasmonic acid treatment. These proteins have a catalytic triad consisting of Ser (dark cyan arrow), His (purple arrow), and Asp (red arrow). Annotations are otherwise as in **Supplementary Figure S1**.

**Figure S16:**
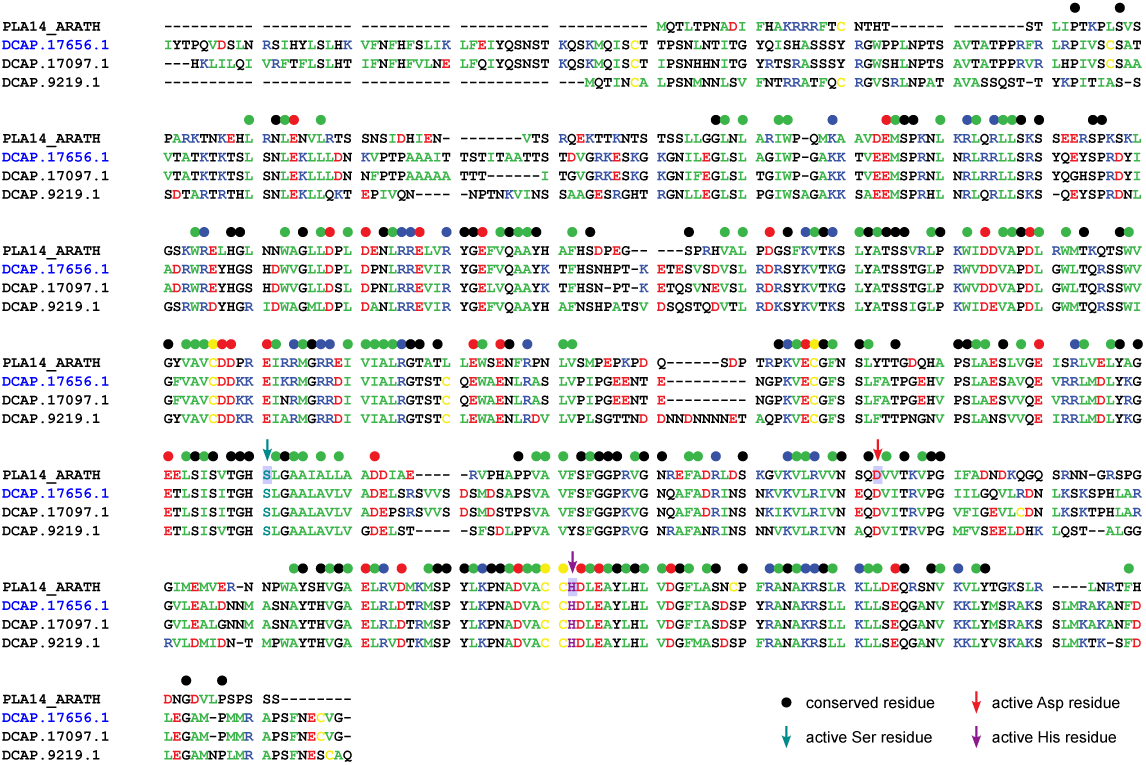
Phospholipase A1 homologs. *D. capensis* has three homologs of *Arabidopsis* phospholipase A1, which hydrolyzes phosphatidylcholine (UniProt ID: PLA14_ARATH) (*138*). One is downregulated in response to jasmonic acid treatment. These proteins have a catalytic triad consisting of Ser (dark cyan arrow), His (purple arrow), and Asp (red arrow). Annotations are otherwise as in **Supplementary Figure S1**.

**Figure S17:**
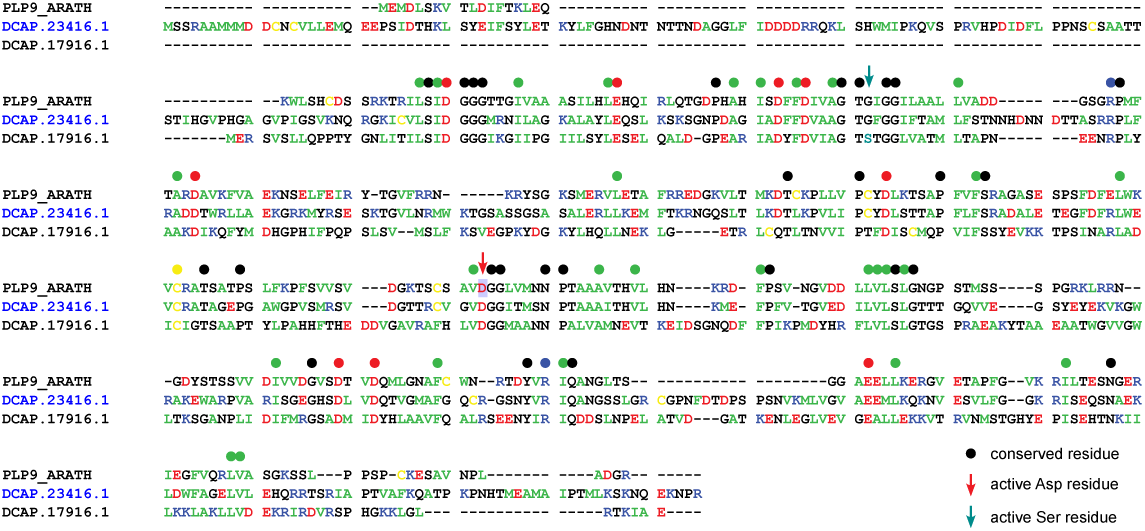
Probable inactive patatin-like protein 9 homologs. *D. capensis* has two homologs of *Arabidopsis* Probable inactive patatin-like protein 9 (UniProt ID: PLP9_ARATH). One is upregulated in response to jasmonic acid treatment. PLP9 is missing the catalytic Ser residue, so its function is unknown. The upregulated protein is likewise missing the catalytic Ser, but it is present in the one that is unchanged in response to jasmonic acid (dark cyan arrow). Annotations are otherwise as in **Supplementary Figure S1**.

**Figure S18:**
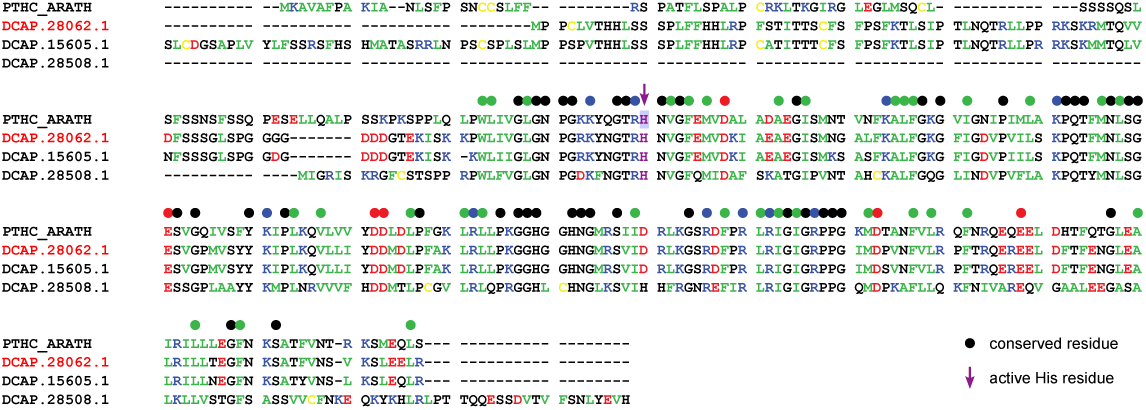
Peptidyl tRNA hydrolase. *D. capensis* has three homologs of *Arabidopsis* Peptidyl-tRNA hydrolase (UniProt ID: PTHC_ARATH). One of these is upregulated in response to jasmonic acid treatment. The active residue is a His (purple arrow). Annotations are otherwise as in **Supplementary Figure S1**.

**Figure S19:**
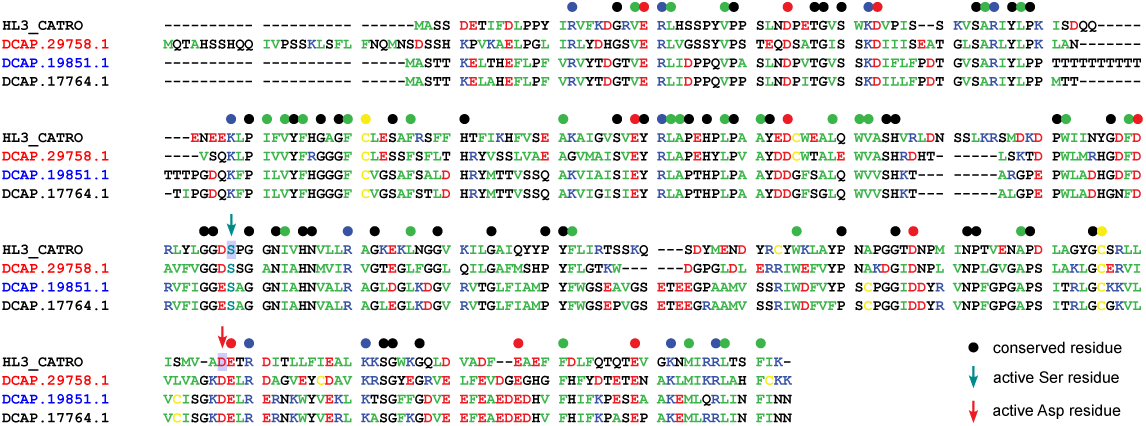
Hydrolase 3 homologs. *D. capensis* has three homologs of *Catharanthus roseus* hydrolase 3 (UniProt ID: HL3_CATRO), which is involved in the biosynthesis of secondary metabolites (*139*). One of these is upregulated and one downregulated in response to jasmonic acid treatment. These proteins have a catalytic dyad consisting of Ser (dark cyan arrow) and Asp (red arrow). Annotations are otherwise as in **Supplementary Figure S1**.

**Figure S20:**
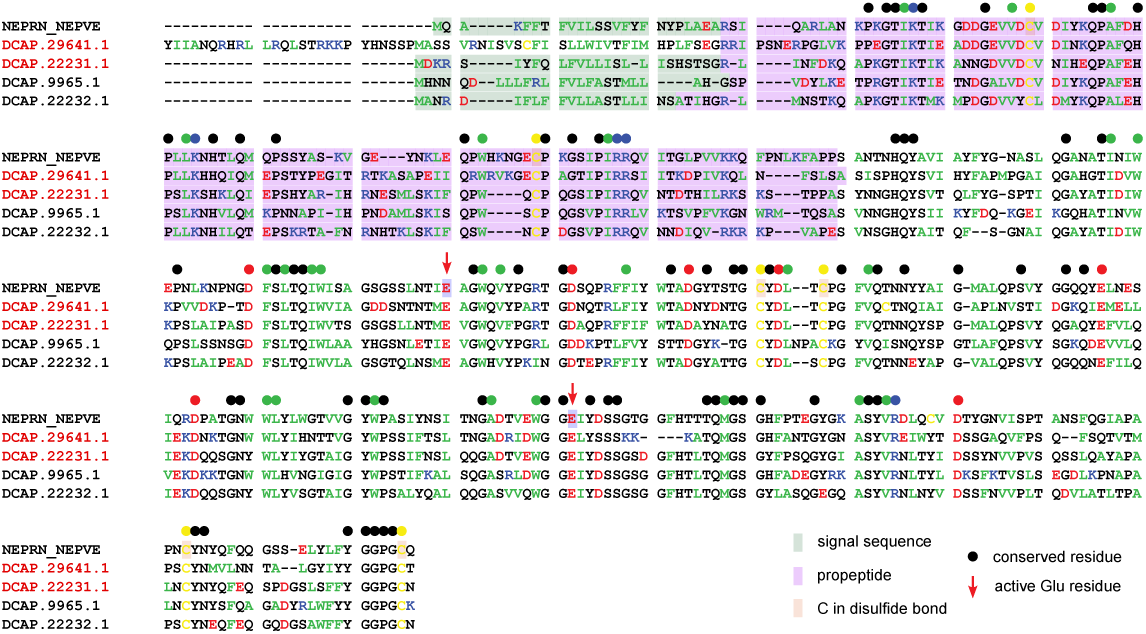
Neprosins. *D. capensis* has 4 neprosins, which are glutamic proteases. Two are upregulated in response to jasmonic acid, while two others remain unchanged. They are annotated by homology to neprosin from *Nepenthes* x ventrata (UniProt ID: NEPRN_NEPVE). The active glutamic acid residues are indicated with red arrows. Annotations are as in **Supplementary Figure S1**.

**Figure S21:**
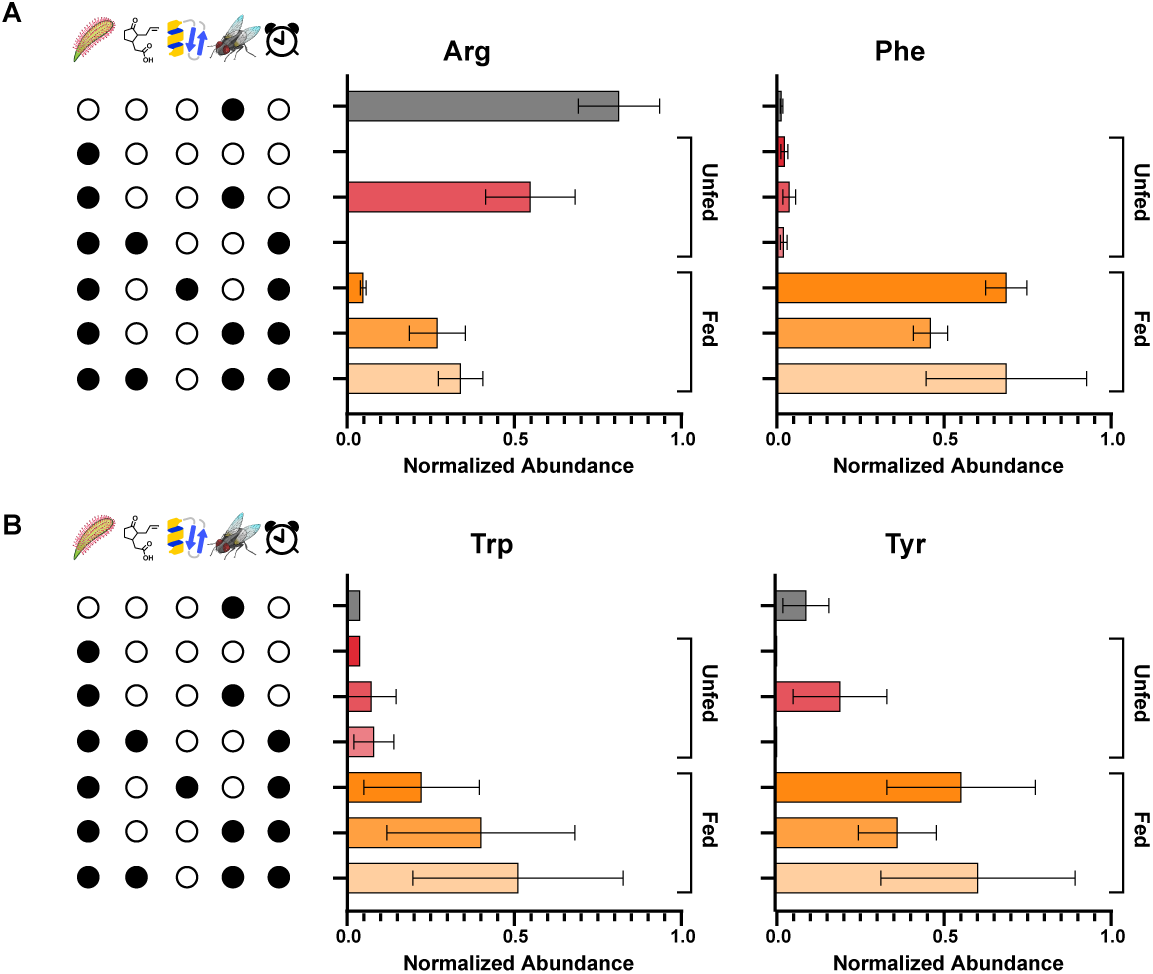
Intensities of amino acids detected for *D. capensis* leaves under different experimental conditions. Individual bar plots showing the average relative abundance of each amino acid in **Figure 5**, in response to each treatment. Bars show mean relative abundance, calculated from the relative abundance of all replicates in the treatment group across features, ± 95% confidence interval.

**Figure S22:**
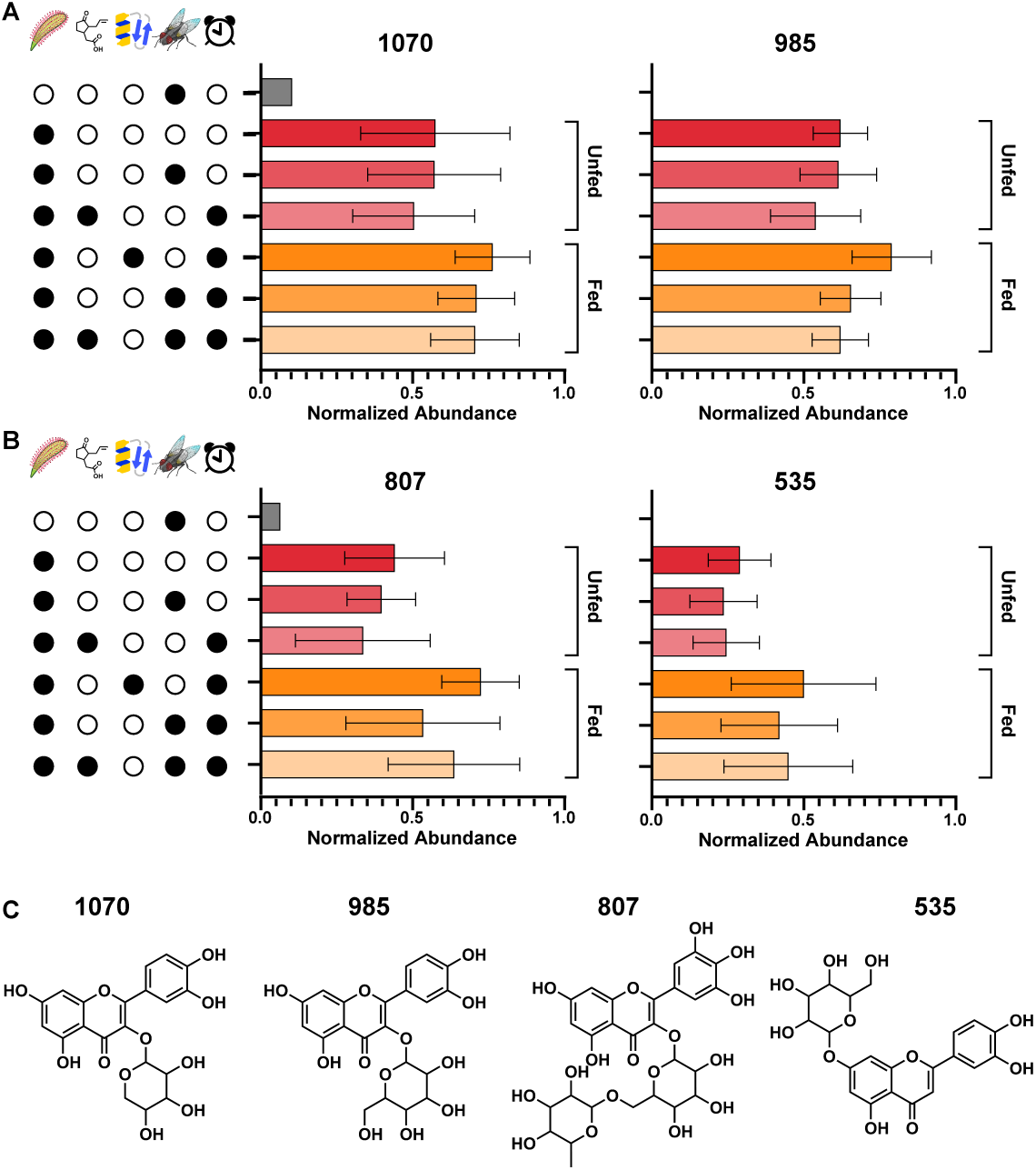
Intensities of flavonoids and anthocyanins detected for *D. capensis* leaves under different experimental conditions. Same as **Supplementary Figure S21**, but for **Figure 6**, showing flavonoid abundances.

**Figure S23:**
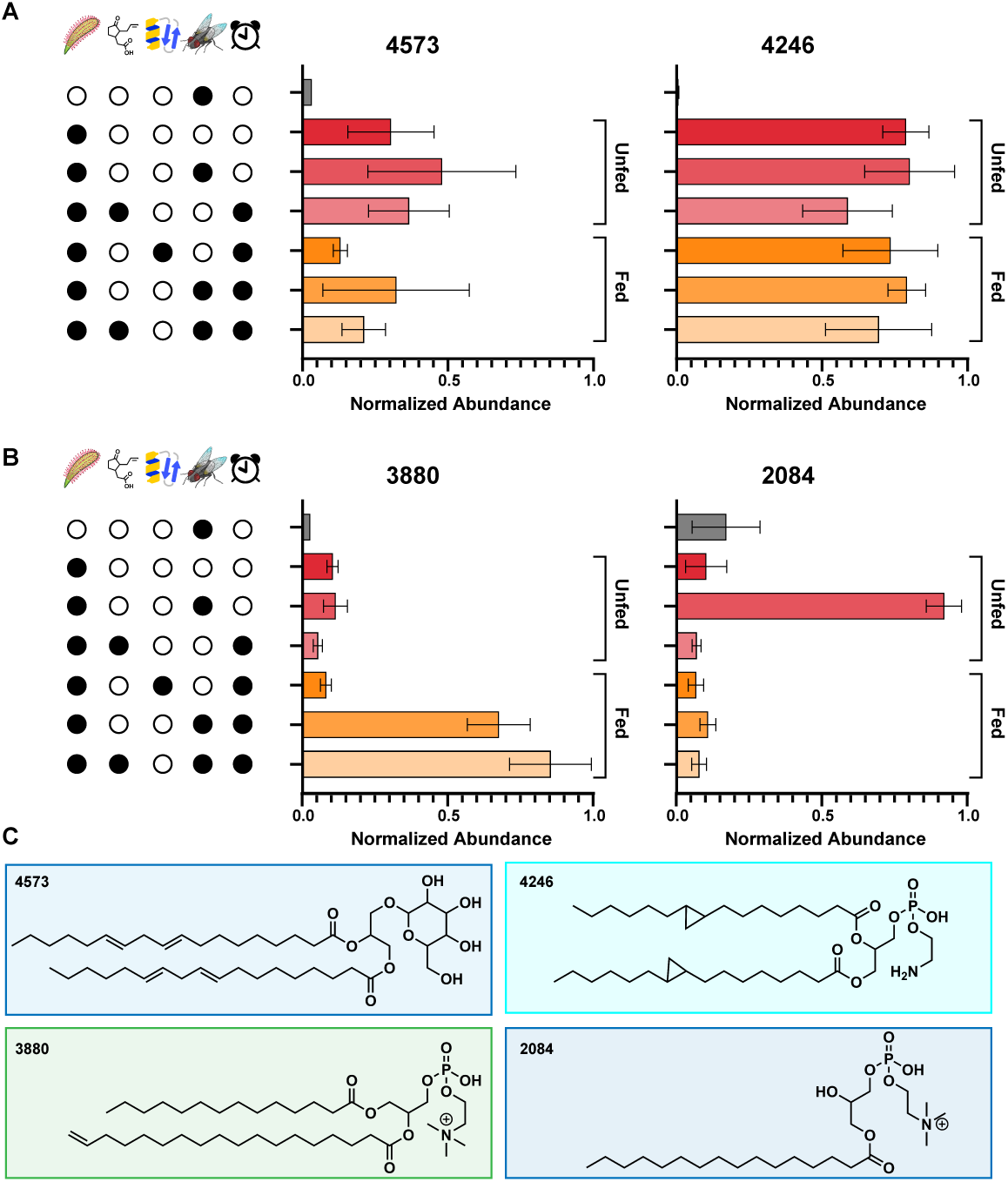
Intensities of lipids detected for *D. capensis* leaves under different experimental conditions. Same as **Supplementary Figure S21**, but for **Figure 7**.

**Figure S24:**
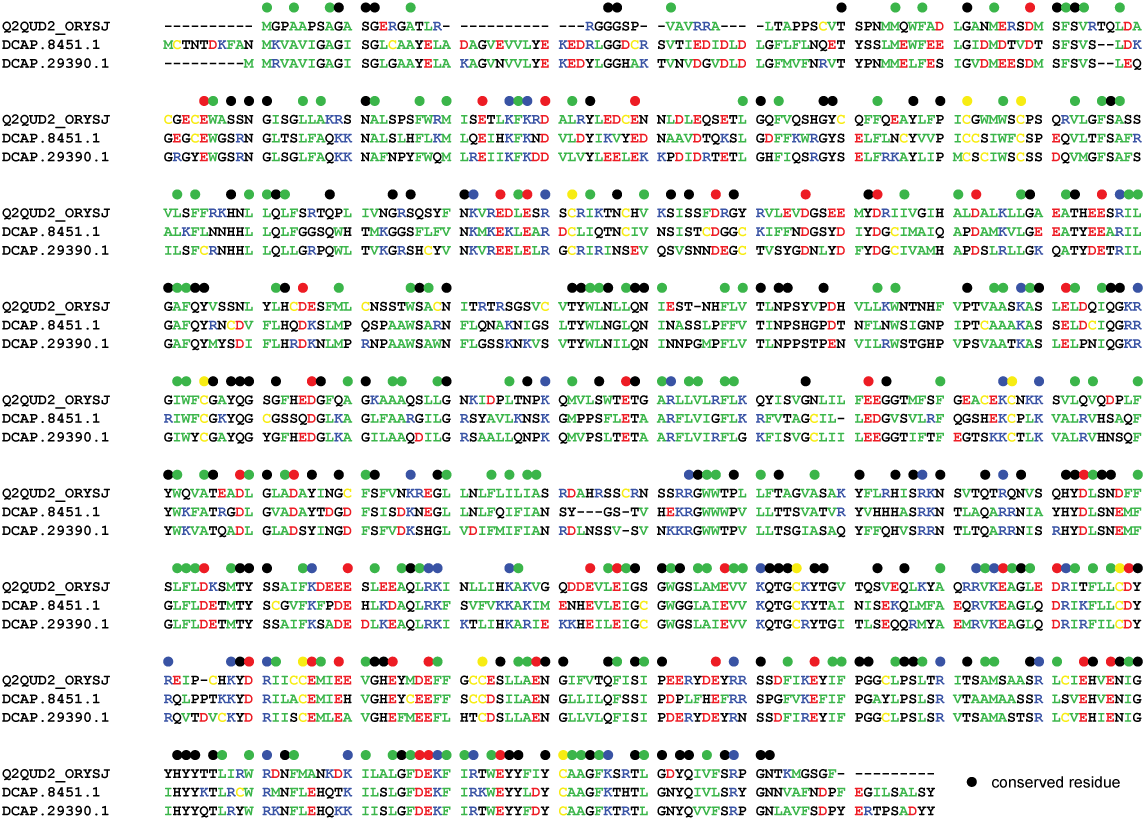
Cyclopropane fatty acid synthases. These proteins align well with Q2QUD2_ORYSJ, a putative cyclopropane fatty acid synthase from rice (*Oryza sativa* subsp. japonica). Annotations are as in **Supplementary Figure S1**.

**Figure S25:**
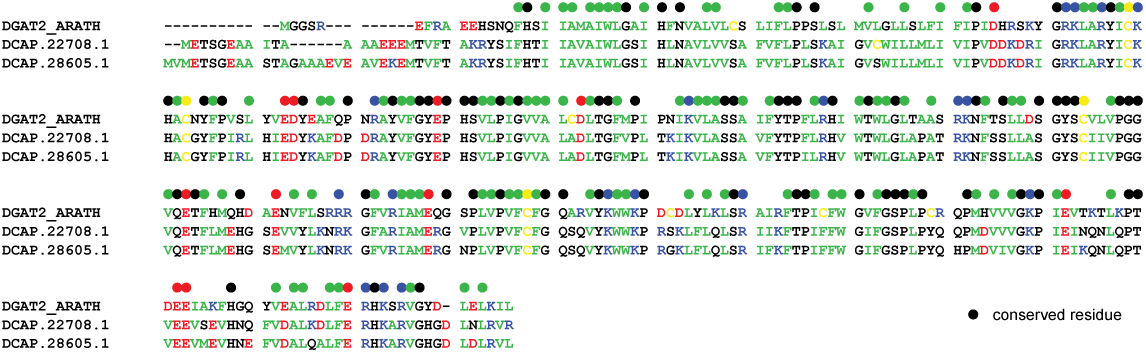
Diacylglycerol O-acyltransferase 2 paralogs. These proteins align well with diacylglycerol O-acyltransferase 2 (UniProt ID: DGAT2_ARATH) from *Arabidopsis thaliana* (*87*). Annotations are as in **Supplementary Figure S1**.

**Figure S26:**
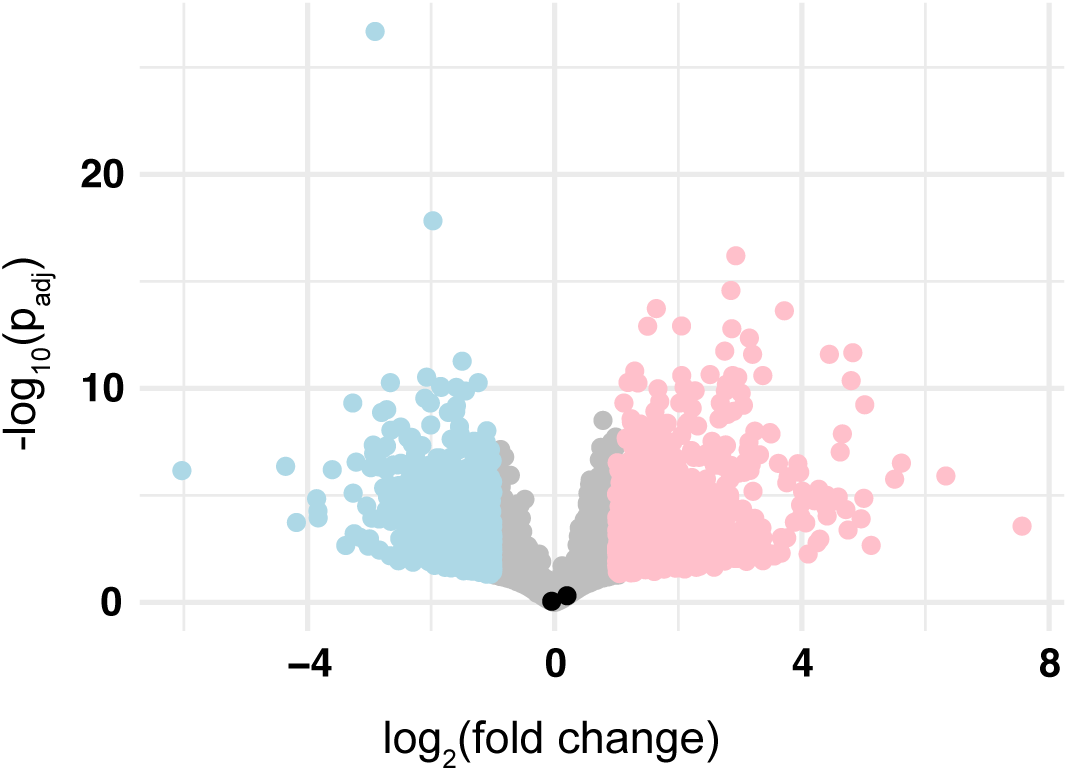
Volcano plot showing the lack of up- or down-regulation of cyclopropane synthases in response to jasmonic acid treatment. *Drosera capensis* has two cyclopropane synthase paralogs; neither changes its expression level in response to jasmonic acid treatment.

**Figure S27:**
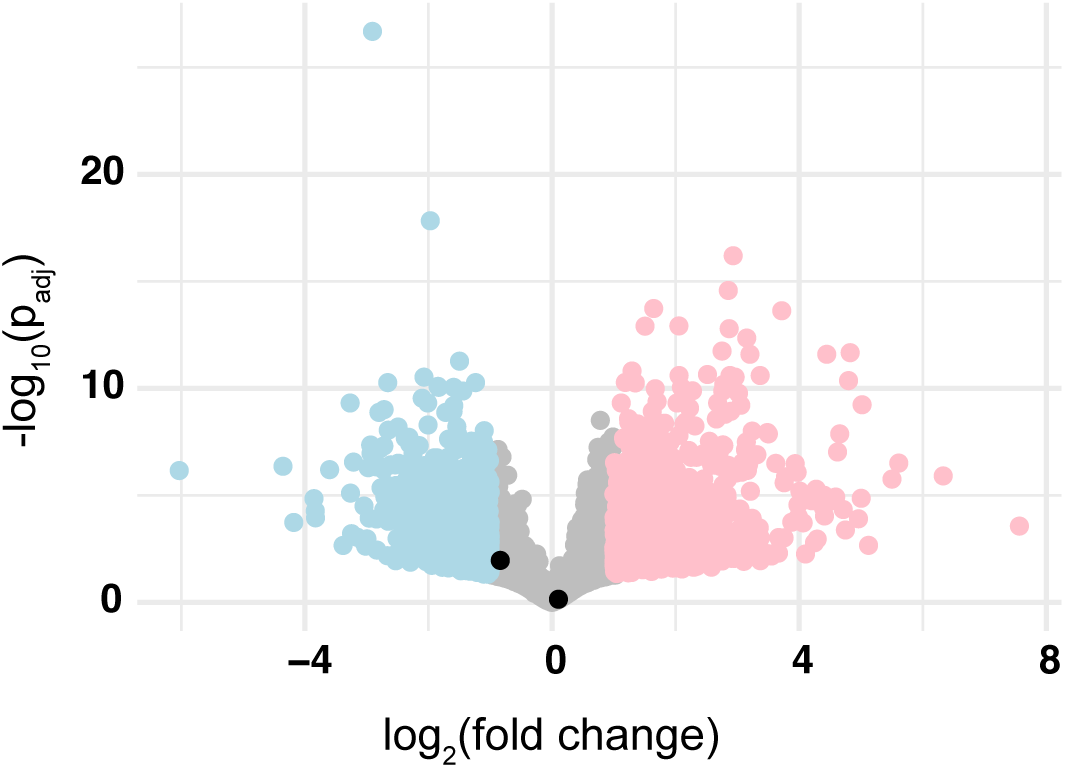
Volcano plot showing the lack of up- or down-regulation of DGAT2 in response to jasmonic acid treatment. *Drosera capensis* has two DGAT2 paralogs; neither changes its expression level in response to jasmonic acid treatment.

**Figure S28:**
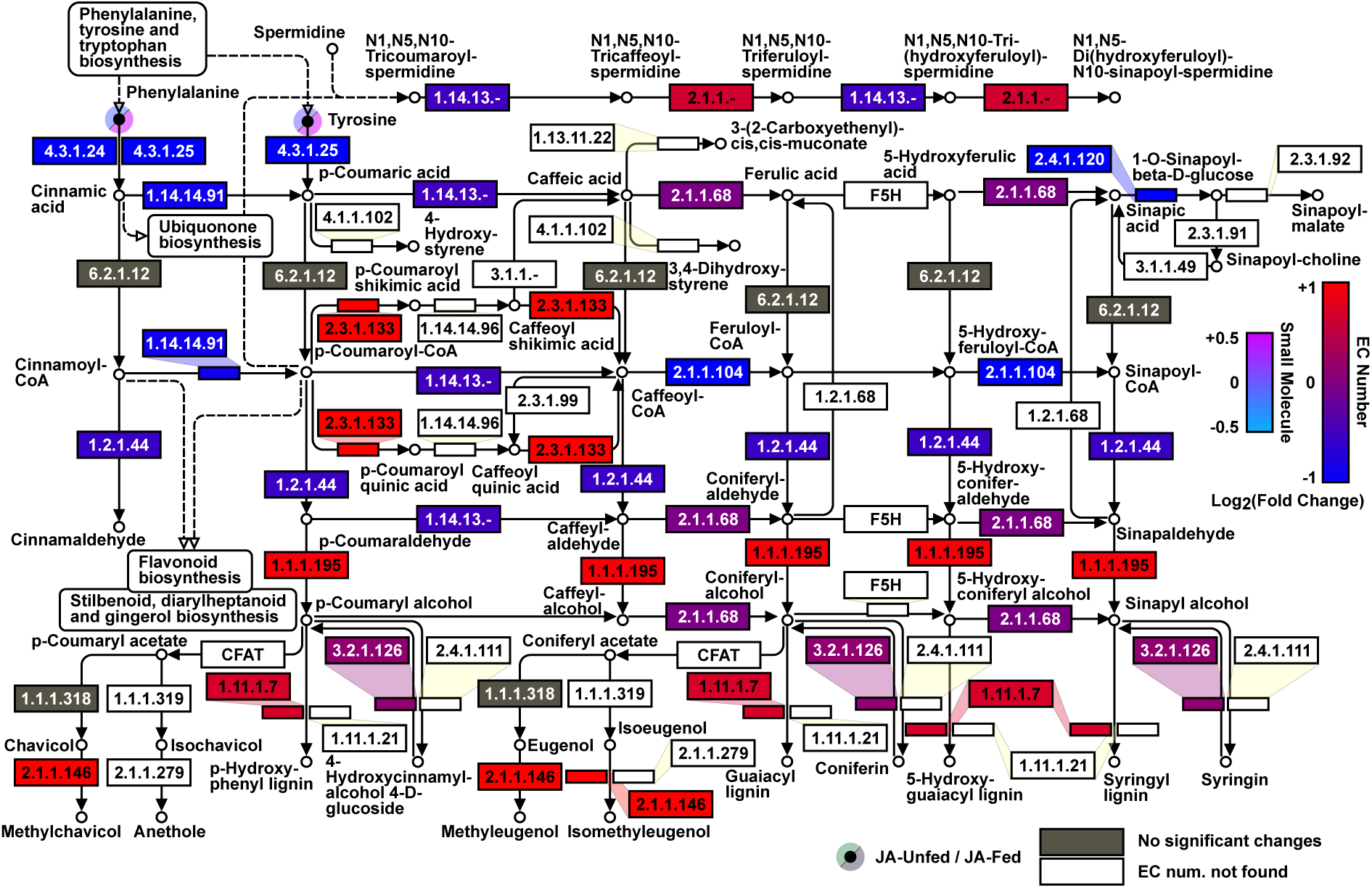
Differential expression of enzymes and small molecules in the KEGG phenylpropanoid biosynthesis pathway (*71–73*) upon treatment with jasmonic acid (JA-Unfed) or jasmonic acid and BWH (JA-Fed). Only genes with padj*<*0.05 (significant genes) contributed to the blue/red coloration in this figure. The blue/red color is the average *log*_2_FC of all significant genes with assigned EC numbers. Color thresholds (−1 and +1 for enzyme *log*_2_FC, −0.5 and +0.5 for small molecule *log*_2_FC) do not indicate the true lower and upper limits of *log*_2_FC for these categories; some values exceed these thresholds. Phenylalanine and tyrosine *log*_2_FC during the JA-Fed condition, which far exceeded the +0.5 threshold that was set, can be visualized in **Figure S21**. EC numbers in grey were found in the transcriptome, but were not assigned to any significant genes. EC numbers in white were not found in the transcriptome.

**Data File S1. Amino acids and amino acid derivatives found using Sirius** Annotations for the amino acids shown in **Figures 5** and other amino acid derivatives are provided in the Supplementary Data File AminoAcids.csv. The data columns are as follows: (A) Sirius Feature ID - ID number output by the Sirius software (B) Feature m/z in Daltons/charge unit (C) Feature retention time in minutes (D) Compound Name - common name or IUPAC name (E) Adduct - adduct observed in the positive mode experiment, generally corresponding to the addition of either H^+^ or Na^+^. (F) CSIFingerID Score - a ‘quality score’ generated by Sirius to describe how well the data overall fit the structure (G) Tanimoto Similarity - a quality score describing how well the MS2 spectra match the theoretically expected spectra (H) Molecular Formula (I) InChI Key - International Chemical Identifier (J) SMILES - representation of the molecular formula for each compound using the Simplified Molecular Input Line Entry System, which describes chemical structures using short ASCII strings. (K) Min - the minimum signal intensity for each feature in arbitrary units (L) Max - the maximum signal intensity for each feature in arbitrary units (M) Difference - the difference between the maximum and minimum intensity for each feature (N) FC - fold change (O) Highest Group - the experimental group for which the maximum signal intensity occurs.

**Data File S2. Flavonoids and anthocyanins found using Sirius** Annotations for the flavonoids shown in **Figures 6A** are provided in the Supplementary Data File Flavonoids.csv. Columns are the same as for Supplementary Data File S1.

**Data File S3. Lipids found using Sirius** Annotations for the lipids shown in (**Figures 7A** are provided in the Supplementary Data File Lipids.csv. Columns are the same as for Supplementary Data File S1, with the addition of column (P) Compound Class.

**Data File S4. Small molecules belonging to the flavonoid pathway** List of molecules referred to by number in (**Figures 8B** are provided in the Supplementary Data File Flavonoid_Pathway_ID. Data columns are as follows: (A) - number (B) - molecule name (C) SMILES code

